# Mutational signatures of replication timing and epigenetic modification persist through the global divergence of mutation spectra across the great ape phylogeny

**DOI:** 10.1101/805598

**Authors:** Michael E. Goldberg, Kelley Harris

## Abstract

Great ape clades exhibit variation in the relative mutation rates of different three-base-pair genomic motifs, with closely related species having more similar mutation spectra than distantly related species. This pattern cannot be explained by classical demographic or selective forces, but imply that DNA replication fidelity has been perturbed in different ways on each branch of the great ape phylogeny. Here, we use whole-genome variation from 88 great apes to investigate whether these species’ mutation spectra are broadly differentiated across the entire genome, or whether mutation spectrum differences are driven by DNA compartments that have particular functional features or chromatin states. We perform principal component analysis and mutational signature deconvolution on mutation spectra ascertained from compartments defined by features including replication timing and ancient repeat content, finding evidence for consistent species-specific mutational signatures that do not depend on which functional compartments the spectra are ascertained from. At the same time, we find that many compartments have their own characteristic mutational signatures that appear stable across the great ape phylogeny. For example, in a mutation spectrum PCA compartmentalized by replication timing, the second PC explaining 21.2% of variation separates all species’ late-replicating regions from their early-replicating regions. Our results suggest that great ape mutation spectrum evolution is not driven by epigenetic changes that modify mutation rates in specific genomic regions, but instead by trans-acting mutational modifiers that affect mutagenesis across the whole genome fairly uniformly.

**SIGNIFICANCE STATEMENT:** All heritable variation begins with damage or copying mistakes affecting the DNA of sperm, eggs, or embryos. Different DNA motifs can have different mutation rates, and these rates can evolve over time: the spectrum of mutability of three-base-pair motifs has evolved rapidly during great ape diversification. Here, we show that even as ape mutation spectra diverged from each other, ape genomes preserved a landscape of spatial mutation spectrum variation. We can thus deconvolute the mutational process into a mixture of fast-evolving signatures with uniform spatial distributions and conserved signatures that target specific regions. Our findings may ultimately help determine the factors, either genetic or environmental, that contribute to temporal and spatial variation in germline mutagenesis.

## INTRODUCTION

The pace of evolution and the healthspan of somatic tissue are both ultimately limited by the genomic mutation rate, which is a complex function of DNA damage susceptibility, polymerase fidelity, proofreading efficacy, and other factors (Alexandrov et al. 2013; Sima & Gilbert 2014). Some regions of the genome accumulate mutations faster than others, such as DNA that replicates late in the cell cycle and motifs with certain epigenetic modifications (Koren et al. 2012; Liu et al. 2013; Polak et al. 2015; Schuster-Böckler & Lehner 2012; Sima & Gilbert 2014; Stamatoyannopoulos et al. 2009). Such mutation rate differences have the potential to confound efforts to infer patterns of purifying selection and background selection (Hellmann et al. 2005; Martincorena et al. 2012). As a result, understanding how mutation rate varies across the genome is an important prerequisite for identifying modes and targets of natural selection (Hellmann et al. 2003; Keightley et al. 2011; Kulathinal et al. 2008). Understanding the mutational landscape is similarly essential for predicting rates of deleterious *de novo* mutations in clinically relevant disease genes (Michaelson et al. 2012; Veltman & Brunner 2012).

Some variation of mutation rate across the genome appears to be driven by features such as chromatin state, transcription, or, more broadly, genomic function (Ananda et al. 2011; Li & Luscombe 2018). For the purpose of this manuscript, we broadly consider a genomic compartment’s “function” to be a set of shared molecular interactions that might or might not be critical to organismal fitness. However, even with this broad definition, a large component of the variance of the mutational landscape still escapes association with any known motifs or functions and has thus been classified as “cryptic” (Hodgkinson & Eyre-Walker 2011; Johnson & Hellmann 2011; Terekhanova et al. 2017). This cryptic variation is conserved over relatively long timescales, being highly similar between human and macaque, which diverged ∼25mya (Tyekucheva et al. 2008). This conservation suggests that many regions of the genome have intrinsic or epigenetic features that affect their mutability and are functionally important enough to be maintained over time despite creating excess deleterious mutation load.

One powerful tool for disentangling the effects of selection and mutation rate variation is mutation spectrum analysis: a comparison of the relative abundance of specific types of mutations, often defined by their triplet context (i.e. AAA>ACA or ACG>ATG) (Hwang & Green 2004). When background selection removes genetic variation from a genomic region, it removes variants that are essentially sampled at random from the spectrum of variation that is present. Biased gene conversion can affect the mutation spectrum, but only in a specific way, favoring the retention of mutations from A/T to G/C over mutations from G/C to A/T (Galtier et al. 2001). A much broader variety of mutation spectrum changes can occur when the mutation rate increases as a result of alteration to the mechanisms of DNA damage or repair, most famously in cancer where cells’ housekeeping processes often break down (Alexandrov et al. 2013). For example, tumors that replicate their DNA with a defective polymerase ε accumulate high rates of TCT>TAT and TCG>TTG mutations (Alexandrov et al. 2013; Shinbrot et al. 2014). Similar “mutational signatures” also occur in the normal human germline, where late-replicating DNA consistently accumulates proportionally more C>A and A>T mutations compared to DNA that replicates earlier during the cell cycle (Agarwal & Przeworski 2019).

In addition to varying between regions of the genome, mutation rates and spectra also vary between different evolutionary lineages. Patterns of diversity point to a global mutation rate slowdown during hominoid evolution that has caused humans and closely related apes to accumulate mutations more slowly than distantly related monkeys do; this slowdown has been studied since the 1980s (Goodman 1985; Scally & Durbin 2012; Li & Tanimura 1987). A closer examination of ape mutation spectra recently revealed that every ape lineage has experienced changes in the relative mutation rates of some characteristic triplet motifs (Harris & Pritchard 2017). Even more surprisingly, closely related human populations have distinctive mutation spectra that provide enough information to classify individuals into continental ancestry groups (Harris & Pritchard 2017). Sometime during the 100,000 years since their descendants migrated out of Africa, Europeans experienced a temporary pulse of mutagenic activity that more than doubled the rate of TCC>TTC mutations (Harris 2015; Speidel et al. 2019; DeWitt et al. 2020).

Mutation spectrum divergence has the potential to shed light on both the mechanisms and overall speed of mutation rate evolution. In theory, many different biological mechanisms can cause mutation rates to evolve; these may be changes to cellular machinery (e.g., changes to a DNA repair protein) or to environmental/life history traits (e.g., longer generation time) (Gao et al. 2019; Shinbrot et al. 2014). In the event that all mutagenic processes generate diversity in a conserved and clocklike manner, mutation spectra are expected to stay constant over time. Conversely, if exposure to a particular mutagen goes up or a DNA repair mechanism becomes less efficient, this is expected to elevate the relative dosage of mutations in genomic motifs that are most vulnerable to damage by that mutagen or preferentially targeted by that DNA repair mechanism. Different genetic and environmental perturbations might cause similar changes in the overall mutation rate, but are less likely to elevate mutation rates in the same genomic sequence contexts.

In this study, we examine how mutation spectra covary across the genome and the ape phylogeny and find that great ape mutation spectra exhibit similar patterns of mutation spectrum divergence across both slowly and quickly mutating compartments of the genome. We find no evidence that species-specific mutator activity is correlated with chromatin state, ancient repeat content, or replication timing during S phase, despite the fact that all of these variables correlate with stable mutational signatures that are consistently detectable across the great ape phylogeny. This implies that the rapid evolution of great ape mutation spectra has likely been driven by *trans*-acting mutators that do not preferentially target any specific genomic compartments, at least not compartments that are correlated with the variables examined in this study. Although there exist many differences between functionally divergent compartments in the spectra of mutations that accumulate, these differences appear to exhibit considerable stability between great ape species and are not likely responsible for the rapid mutation spectrum divergence. We find that such stability extends to species as divergent as humans and mice, where we find consistent mutational signatures present in genomic regions that are not homologous but are annotated with the same functional states as promoters, enhancers, or repressed regions.

## RESULTS

### Quantifying the Mutation Spectrum Differences Between Great Ape Species and Subspecies

Previous research utilizing the Great Ape Genome Project (GAGP) data showed that the germline mutation spectrum has evolved rapidly in great apes, leading to distinct species-specific spectra (Harris & Pritchard 2017; Prado-Martinez et al. 2013). We first sought to recapitulate these results and measure for the first time how the differences between species compare to differences within species (Table S1).

To minimize the effects of natural selection and read mapping errors on our mutation spectrum ascertainment, we defined a set of genomic regions, collectively called a “compartment”, characterized as non-conserved and non-repetitive (NCNR). This NCNR compartment consists of 1.28Gb of the non-repetitive (annotated by RepeatMasker), non-coding human genome excluding both significantly conserved regions (*p*<0.05 in the PhastCons 44-way primate alignment) and CpG islands (Table S2). In these compartments, we computed the relative abundances of each of the 96 triplet mutation types for each individual and each species (Figure 1A) using SNVs from the GAGP, following a number of filters. To ensure that shared genetic drift cannot inflate the appearance of mutation spectrum similarity among individuals that share many derived alleles, we computed mutation spectra using a sampling procedure that randomly counts each SNV toward the spectrum of only one of the individuals that carry it, rather than all such individuals (see Methods). This sampling method reduces the impact of GC-biased gene conversion, which primarily affects higher frequency alleles in regions specific to lineages (see Methods, Figure S1).

**Figure 1:**
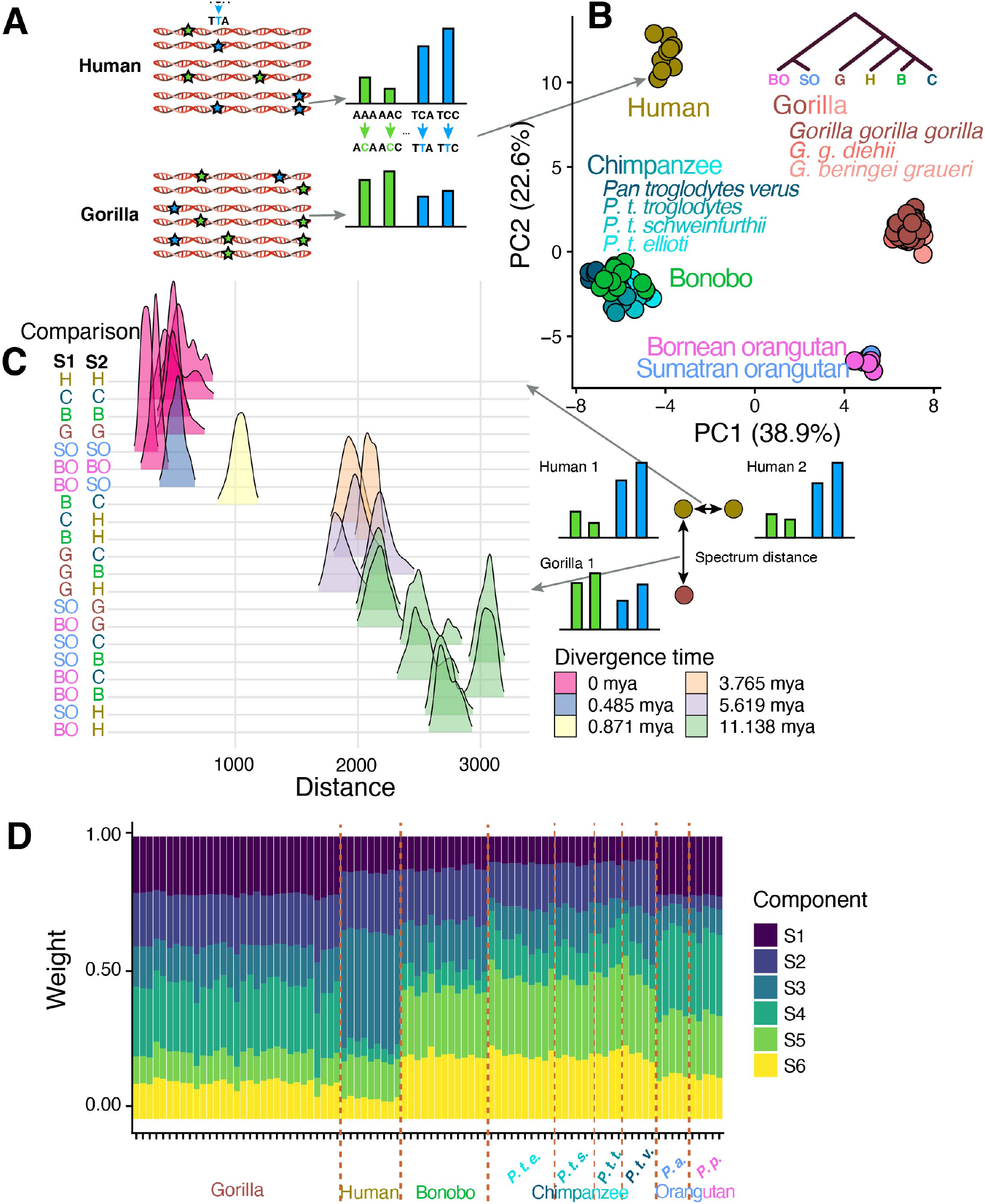
Covariance of species-specific and replication timing mutation spectra in great apes. A. SNVs segregating within a species were counted to generate a triplet mutation spectrum for each individual in the GAGP. We include SNVs found in non-conserved, non-repetitive (NCNR) regions of the genome, which we collectively call the NCNR compartment. B. PCA of NCNR compartment mutation spectra reveals clustering of individuals by species. Each point represents the NCNR compartment mutation spectrum from a single individual in the GAGP; colors represent species, while shades of a color represent subspecies (applies only to gorillas and chimpanzees). C. Mutation spectra are more similar between individuals of the same species than between individuals from different species. We plotted the Euclidean distances between triplet mutation spectra of all possible pairs of individuals in the GAGP within and between species (see Methods). D. When nonnegative matrix factorization (NMF) is used to infer mutational signatures that best explain variation among ape mutation spectra, we see more variation of signature composition between species than within species and identify signatures that change in dosage on specific phylogenetic tree branches. For example, signature 1 appears to have decreased in dosage on the branch ancestral to humans, chimps, and bonobos.

A principal component analysis (PCA) on mutation spectra shows clustering of individuals by species in a manner that recapitulates phylogeny. This pattern reveals the existence of distinct species-specific mutation spectra (Figure 1B). Mutation spectra of humans, chimpanzees (*Pan troglodytes*), and bonobos (*Pan paniscus*), which form a phylogenetic clade, separate from more distantly related gorillas (*Gorilla gorilla*) and Sumatran and Bornean orangutans (*Pongo abelii* and *Pongo pygmaeus*, respectively) along principal component 1 (PC1). PC2 separates humans from chimpanzees and bonobos in addition to separating gorillas from orangutans. The two orangutan species cluster closely together, which is unsurprising given that their divergence time (∼483kya) is an order of magnitude more recent than that of humans and chimpanzees (Prado-Martinez et al. 2013). PC1 appears dominated by the proportion of A>C, and A>T and components of C>T mutations, while PC2 is dominated by the proportion of C>A and C>G mutations (Figure S2). The major A>C component of PC1, for example, corresponds to a 10-15% decrease in the fraction of A>C SNVs in gorillas and orangutans relative to humans, chimpanzees, and bonobos. PC2’s C>A component corresponds to a 4% and 7% increase in the C>A SNV fraction in humans and gorillas relative to the *Pan* and *Pongo* clades, respectively. These results are robust to subsampling of equal numbers of individuals across species (Figure S3). Although prior papers have reported homogeneity of de novo mutation spectra between non-human great apes, we find that these studies are underpowered to detect species differences of the magnitude we observe here (Thomas et al. 2018; Besenbacher et al. 2019) (See Methods, Table S3).

We even observed mutation spectrum differences among chimpanzee subspecies, as visible in the PCA: Western chimpanzees (*P. troglodytes verus*) overlap slightly less with bonobos along PC1 than other chimpanzee subspecies, which begin to separate from bonobos along PC2. The mutation spectrum differences between Western and non-Western chimpanzee subspecies is more clearly visualized in a PCA run on those individuals alone (Figure S4). The first PC demonstrates that the variance in mutation spectra between Western and non-Western chimpanzees significantly exceeds the variance among non-Western chimpanzees (p ≤ 2.2 * 10^−16^, two-sided t-test). Western chimpanzees experienced a population bottleneck when they diverged from the lineage ancestral to other chimpanzees, which might have accelerated their mutation spectrum divergence by allowing mutator alleles to drift to higher frequency (Prado-Martinez et al. 2013; Sudmant et al. 2013; Lynch 2008; Lynch et al. 2016; Kimura 1967) (Figure S4). Gorilla subspecies, orangutan species, and human populations exhibit more subtle mutation spectrum differences that are not visible when spectra are projected onto the principal axes of ape variation (Harris & Pritchard 2017) (Figure S4). To quantify these mutation spectrum differences further, we embedded mutation spectrum histograms as points in a 96-dimensional space and computed distances between the using a standard Euclidean metric. As seen qualitatively in the PCA, we found that interspecific differences exceeded conspecific differences (Figure 1C). Furthermore, interspecific differences scaled with divergence time.

We undertook additional analyses to verify that the observed mutation spectrum differences are not likely caused by the tendency of natural selection to retain variation in certain sequence contexts. Although GC-biased gene conversion does favor the fixation of G/C alleles and the elimination of A/T alleles, none of the mutational signatures that vary in dosage between great ape species are consistent with the spectra of mutations that for which biased gene conversion selects (Figure S2). Similarly, biased gene conversion has similar bias across all species (Figure S5). Furthermore, any mutation spectrum difference caused by the action of natural selection should affect high frequency alleles more than low frequency alleles, so we repeated several key analyses using only low frequency variants. We performed these replicate analyses using doubletons rather than singletons, since singletons are more vulnerable to confounding by sequencing error, and found no qualitative differences from the results obtained using the full frequency spectrum of genetic variants (Figure S6A-B).

We used non-negative matrix factorization (NMF) to explicitly infer which mutational signatures have changed in dosage along different branches of the ape phylogeny (Figure 1D). Similar to PCA, NMF is a model-free method used to determine major components of variance that underlie a matrix of data; the components NMF extracts from mutation spectrum matrices can be interpreted as mutation signatures, following the work of Alexandrov et al., 2013. We ran NMF on the matrix of individual NCNR mutation spectra using Helmsman, specifying the model to infer *K* = 6 signatures (Carlson, Li, et al. 2018). Figure 1D shows the dosage of each signature for every individual in the GAGP, grouped by species. Although each signature is present in every individual, the signatures clearly demonstrate lineage-specific dosage. For example, S3 is present at moderate dosage in all non-human species but appears to have increased in relative rate in humans following the divergence of humans from the human-chimpanzee-bonobo common ancestor (Figure S7). S4, conversely, has decreased 2-fold on the branch leading to human-chimpanzee-bonobo; S2 has decreased nearly 3-fold on the branch leading to orangutans. These results support a prior hypothesis that the dosage of one or more mutational signature has changed along each branch in the great ape phylogeny (Harris & Pritchard 2017).

### A mutational signature associated with DNA replication timing is conserved among great apes

Differences in replication timing explain a substantial portion of the variation in somatic and germline mutation rate across the genome (Koren et al. 2012; Stamatoyannopoulos et al. 2009). Compared to regions that replicate early in S phase, late replicating regions tend to have a higher overall mutation rate, and in humans they particularly harbor a higher rate of A>T and C>A mutations (Agarwal & Przeworski 2019). The established correlation between late replication timing and elevated mutation rate implies that replication timing QTLs (rtQTLs) may be examples of *cis*-acting mutation spectrum modifiers, with late-replication alleles acting to increase the load of a late-replicating mutational signature in the surrounding DNA. We analyzed late- and early-replicating compartments of the genome to determine whether replication timing had a similar effect on the mutation spectrum across great apes.

In an attempt to replicate the late-replication signature reported by Agarwal and Przeworski, we defined early and late replication timing compartments to be the earliest- and latest-replicating quartiles of the genome identified by RepliSeq in human lymphoblastoid cell lines (Koren et al. 2012) (Figure 2A, Table S2). We then generated separate mutation spectra from the early-replicating and the late-replicating compartments of each individual genome, normalizing the spectra by the triplet composition of each compartment. We ran a PCA on a matrix containing three mutation spectra derived from each human genome in the GAGP: two derived from early- and late-replicating compartments and the third derived from the NCNR. compartment (Figure 2B, Figure S8) and observed that PC1 separated out these spectra as a function of replication timing. To determine the principal triplet mutation types driving the differences in mutation spectra between compartments, we generated heatmaps of the log odds ratio enrichments of each mutation type occurring in the late versus the early replication timing compartments (Figure 2C, Figure S9). For this analysis, we counted the number of segregating sites within a species to generate a single 96-dimensional vector for each species and compartment (rather than a vector for each individual and compartment, see Methods). As expected, late-replicating regions were enriched for C>A and A>T mutations.

**Figure 2:**
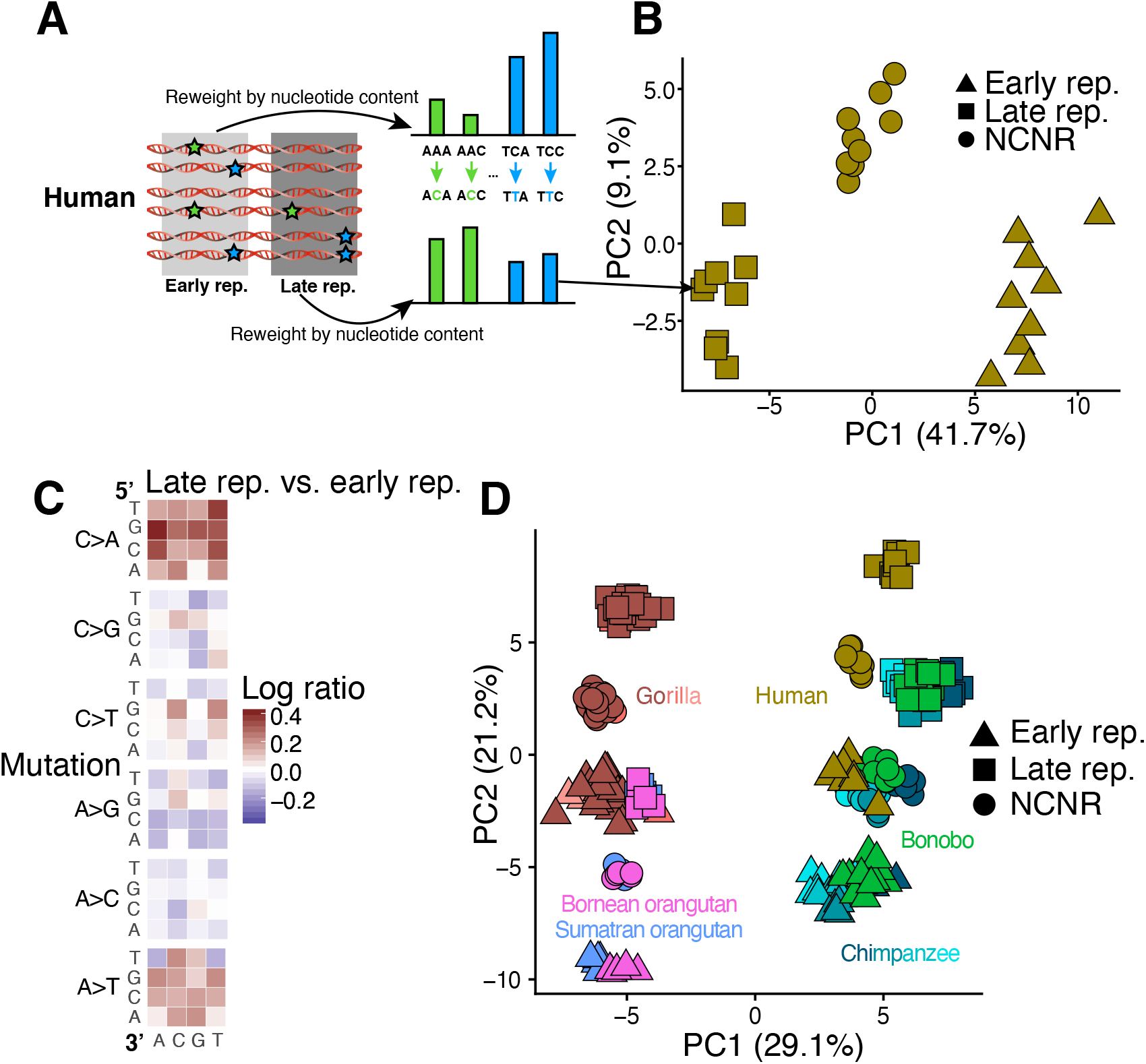
Signatures associated with replication timing appear conserved among great apes A. We calculated separate mutation spectra for each individual in compartments that replicate early versus late in S phase and re-weighted each spectrum by the trinucleotide content of the associated compartment. B. Each point in this PCA represents the mutation spectrum from a single individual’s NCNR, early replicating, or late replicating compartment. The observed gradient results from differences in mutation spectra between early-replicating and late-replicating compartments. C. A heatmap of the log ratios of triplet mutation fractions in humans shows an enrichment for C>A and A>T mutations in the late replicating compartment compared to the early replicating compartment. This mutation signature recapitulates recently described late replication timing signature in humans. To generate the species mutation spectra, we counted the number of SNVs with triplet context segregating within a species that occurred in each compartment. The triplet mutation fractions were normalized by compartment nucleotide content. D. Mutation spectra from late and early replication timing and NCNR compartments from each individual in the GAGP separate along a first PC associated with phylogeny and a second PC associated with replication timing. The mutation signature associated with late replication timing appears conserved among all great apes. Different shades of each species’ color represent subspecies, as in Figure 1D.

After replicating the reported effect of replication timing on the human mutation spectrum, we set out to determine if the action of this signature appeared conserved among species. To this end, we identified the ape genomic compartments that aligned to the human early-replicating and late-replicating compartments and ran a PCA on a matrix of the early replicating, late replicating, and NCNR compartment mutation spectra of all individuals (Figure 2D). PC1 separates the spectra by phylogeny and species identity, while PC2 separates them along an axis that aligns with replication timing. PC3 separates human from chimpanzee and bonobo spectra (not shown). The direction of separation of early and late replication timing compartments is similar across all species, implying the action of a conserved mutational process to a consistent compartment of the genome. The most parsimonious explanation for this pattern is that the replication timing landscape is largely conserved across great apes and that late replication exerts a similar mutagenic effect in all great ape species. As above, we obtained consistent results using only doubletons (Figure S6C,D). A PCA of individual mutation spectra, using only doubletons, recreates similar clustering patterns to those presented in Figure 2. Furthermore, the second PC again captures the mutational signature of late replication timing (Fig S6D). The main effect of restricting to rare variants is that it makes humans appear less displaced from chimps along the axis of differentiation by replication timing. These results suggest that differences between species in biased gene conversion or demographic history have a minimal effect on the observed trends (Figure S6).

To further quantify the conservation of the replication timing mutational signature, we tested the correlation of the log odds of late vs. early replication compartments between each pair of species, thereby quantifying the similarity between each species’ late vs. early replication timing mutational heatmaps. The correlations between every pair of species’ replication timing heatmaps were highly significant (Figure S9, Table S4). Our results show that late replication timing is associated with a conserved mutational signature across great apes. Moreover, these mutational patterns suggest that the genomic landscape of replication timing is broadly conserved across species, biasing all genomes toward C>A and A>T mutations in compartments that are directly orthologous to the human late-replicating compartment (Figure S10). The conservation of this mutation signature contrasts with the rapid evolution of the species signatures. We see a few hints of interactions between replication timing and species identity – for example, CG>GG mutations are depleted in late replication timing regions, and the depletion appears slightly stronger in non-human apes than in humans. However, these nonlinear effects are small and higher resolution sequencing data will be needed to verify whether they are true biological signals.

The layering of rapidly evolving species signatures over conserved replication timing signatures was further supported by NMF decomposition. We ran NMF independently for each species on matrices of individual mutation spectra from the early replicating and late replicating compartments, after normalizing for compartment-specific nucleotide content. Each NMF was run to extract *K* = 7 signatures. Following the methodology of Alexandrov et al. 2013, we then grouped similar signatures by their loadings using hierarchical agglomerative clustering (Figure S11). The clustering of these signatures, even following independently-run NMFs, supports the deep conservation of a mutation signature associated with replication timing that spans the great ape phylogeny. Furthermore, the loadings of these signatures are enriched for C>A and A>T mutation types, once again supporting our presented results.

### All great ape genetic variation appears to be shaped by a conserved landscape of *cis*-acting mutational modifiers

We found that all ape DNA, regardless of replication timing, has a consistent mutation spectrum bias that we call a species-specific signature. Late-replicating regions of chimpanzee genomes contain a mixture of both a chimpanzee-specific signature and the same late-replication signature that is found in human genomes (Figure S9). We see little evidence of any mutational signature unique to late-replicating chimpanzee DNA that is not also found in early-replicating chimpanzee DNA or in late-replicating regions of other ape genomes. Furthermore, we see no evidence that species-specific signatures have a rate or dosage that depends on replication timing.

Using published annotations of the human genome, we defined several more overlapping functional compartments to characterize the phylogenetic distribution of spatially localized mutational signatures. We used RepeatMasker to delineate repetitive vs. non-repetitive DNA and used ENCODE chromHMM output (the intersection of heterochromatic regions in nine cell types) to annotate several types of heterochromatin (Ernst & Kellis 2012). Another compartment we annotated consists of ancient repeats, which have the potential to mutate differently from higher complexity DNA via several mechanisms, including the formation of non-B-DNA secondary structures and the editing activity of antiviral enzymes such as APOBECs (Bacolla et al. 2004; Guiblet et al. 2018; Harris et al. 2002) (Table S2).

We ran a PCA for each species comparing the mutation spectra of all eight compartments and observed similar topologies of compartment separation in each species. In all cases, the vector separating late-replicating from early-replicating compartments is nearly aligned with the first principal component, which explains 23.3% to 34.1% of the total variance. PC2, which explains 8.2%-13.5% of the total variance, is similarly aligned with the separation between repetitive and nonrepetitive compartments. Finally, PC3 (4.0%-8.7% variance explained) separates the ERVs from the other compartments to a much greater extent than either of the first PCs (Figure 3A-F, Figure S12). The similarities of the independent PCAs across all species of great apes imply conservation of the *cis*-regulated mutational signatures associated with repetitive content, methylation, and replication timing. Each compartment shows a similar degree of separation between species, with high degrees of correlation between the positioning of compartments in different species’ PCAs (Table S5).

**Figure 3:**
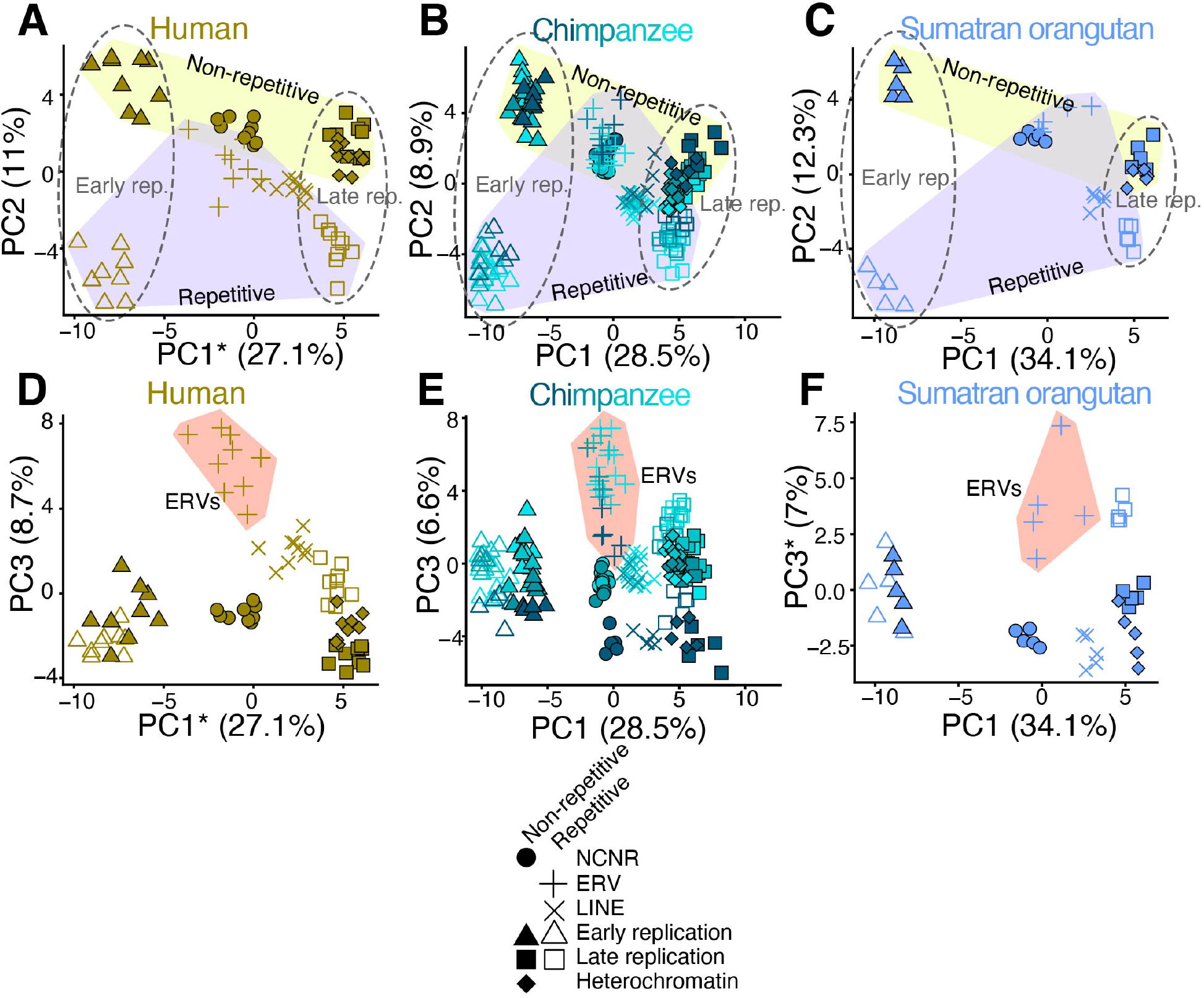
Conserved axes of mutation spectrum variance among great apes A-F. We defined eight overlapping functional compartments to test for evolution of mutation spectrum modifiers along axes of chromatin accessibility, replication timing, and repetitive content. We then ran a PCA on the individual mutation spectra for all eight compartments for each species separately (only human, chimpanzee, and Sumatran orangutan shown). For all species, PC1 and PC2 separate compartments along gradients that correspond to replication timing and repetitive content, respectively (dotted lines vs. shaded polygons, A-C). PC3 separates ERVs from other compartments (shaded polygons, D-F). The similarities of these independent PCAs across all species implies conservation of *cis*-acting mutational signatures. *Axis inverted for readability.

We identified only one genomic compartment whose localized mutational signature appears to be distributed non-uniformly across species: maternal mutation hotspots first identified using human trio data (Jónsson et al. 2017) (Table S2). Maternal mutation hotspots are genomic regions that are enriched for *de novo* C>G mutations that arise on the maternal lineage and whose rate has an unusually strong correlation with maternal age. These hotspots exist in chimpanzees and, to a lesser extent, gorillas, but their signal is nearly absent from orangutans. Our mutation PCAs similarly show that human genomes have higher levels of a compartment-specific mutational signature in these regions (Figure S13). The separation of NCNR to maternal mutation hotspot mutation spectra compartments decays with phylogenetic distance from humans, recapitulating findings from Jonsson et al. (Jónsson et al. 2017). Although a *trans*-acting protein might be involved in the creation of C>G mutations at maternal hotspots, perhaps due to error-prone repair of double strand breaks, the targeting of damage toward specific regions fits the profile of a *cis*-acting targeting factor. Either this *cis* targeting factor or a *trans* interacting partner has been intensifying its mutagenic effects along the evolutionary lineage leading to humans, causing extra mutations to accumulate in a localized pattern. Although differences in maternal age at conception may be a partial explanation for these observations, generation time differences cannot fully explain the patterns across all great apes. The strength of this signature certainly decreases with generation time among humans, *Pan* clade, and gorillas (29, 24, and 19 years respectively).

However, orangutans have a higher average maternal age (25 years) at conception than that of gorillas and the *Pan* clade but exhibit the weakest dosage of the C>G signature (Besenbacher et al. 2019).

The compartment that harbored a signature whose dosage was the strongest among the presently investigated compartments are CpG islands, genomic regions ranging from 200-2000bp long that are enriched for CpG dinucleotides (Gardiner-Garden & Frommer 1987; Larsen et al. 1992) (Table S2). CpG islands are the only compartment we identified whose differentiation from the NCNR compartment explains a greater proportion of mutation spectrum variance than differentiation between species across these regions. Outside of these CpG islands, CpGs are often methylated to 5-methylcytosine, which mutate to TpG at a rate ten times higher than unmethylated CpGs. In contrast, CpG islands are hypomethylated and are often situated in conserved 5’ promoters and genic regions. We hypothesized that, due to their lack of CpG>TpG mutations and overall conservation, a compartment containing CpG islands would demonstrate a contrasting mutation spectrum relative to that of the NCNR compartment. A PCA of individual mutation spectra from the NCNR and a CpG compartment from all GAGP individuals demonstrated that differentiation of the CpG island compartment exceeds the magnitude of spectrum differentiation between species (Figure S14). This is unsurprising given the unique mutational properties of CpG islands compared to the rest of the genome.

### Endogenous retroviruses carry a distinct mutational signature conserved across great apes

ERVs are a class of repetitive, transposable DNA elements that duplicate themselves in a copy and paste manner. The act of duplication into new regions of the genome can disrupt function; for example, integration into the coding region of a gene could result in a complete knock-out of gene function. Therefore, ERV activity is restricted by a number of known mechanisms, including hypermethylation, inhibition of integration, and hypermutation.

In light of these mechanisms that target ERVs, we were intrigued by the fact that ERVs separated from other genomic compartments along the third principal component of our mutation spectrum analysis (4-8% variance explained). ERVs bear an excess load of a unique mutational signature that appears to be largely conserved among great apes (Figure 2B,D,F) and is previously undescribed, to our knowledge.

To determine whether any component of the ERV signature could be caused by high rates of methylation and heterochromatization, we directly compared the mutation spectrum of the ERV compartment to that of nonrepetitive heterochromatin. We calculated the log ratio enrichments and depletions of the 96 mutation types in ERVs relative to nonrepetitive heterochromatin for each species and found an enrichment for CpG C>G mutations and a depletion of TAA>TTA mutations in ERVs that appears conserved in all species other than *Pongo pygmaeus* (Figure 4A). This comparison shows that ERVs’ high rate of CpG>CpT transitions is likely caused by their heterochromatic status, but that heterochromatinization cannot explain the other components of the ERV signature. Furthermore, we determined that differences in 7-mer nucleotide content between the two compartments explained some, but not all, of the ERV-specific enrichment for CG>GG mutation types (Figure S15).

**Figure 4:**
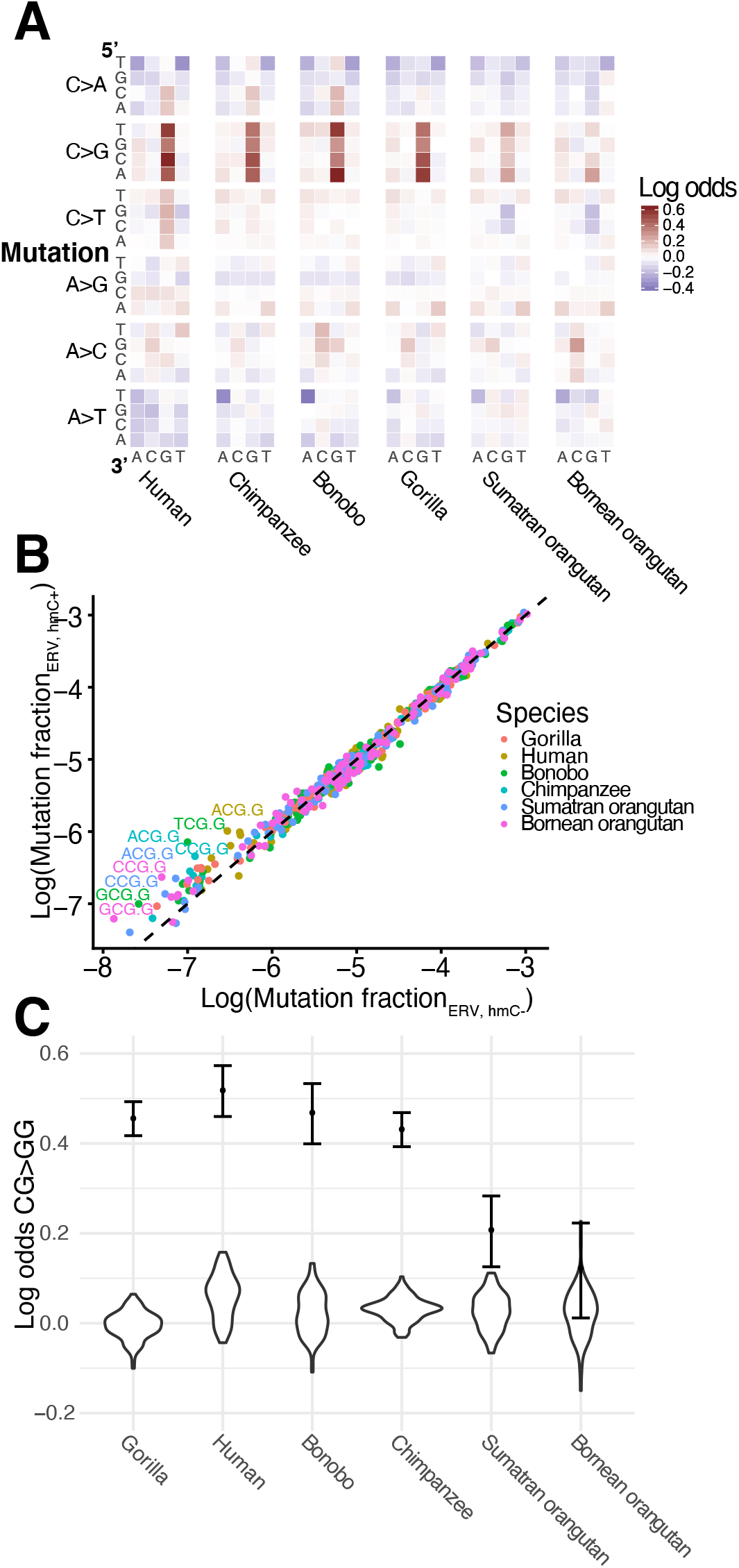
A hydroxymethylation-related CG>GG mutation signature distinguishes ERVs from other compartments A. Heatmaps of log odds of triplet mutation spectra comparing ERV to nonrepetitive heterochromatin compartments for each species show a significant ERV-specific CG>GG mutation signature. B. Enrichment of CG>GG mutation types in ERVs with hydroxymethylation compared to ERVs without. Points represent the fraction of each triplet mutation in ERVs with and without hydroxymethylation calculated from SNVs segregating within each species (*y* and *x* axis, respectively). Mutation types that fall along the *y* = *x* line occur equally frequently in both compartments. Mutation type labels are included only for mutation types whose log ratio of ERV hmC+:ERV hmC-exceeds 0.4. Points are colored by species. C. The CG>GG mutation signature in ERVs is robust to the size and shape of the ERV compartment. We created 100 “ERV-like” compartments by sampling segments corresponding to the size of those in the ERV compartment from random locations within the nonrepetitive heterochromatin compartment. The distribution of the log odds of CG>GG mutations between these ERV-like to the original nonrepetitive heterochromatin compartment are violin plots. The log odds of CG>GG mutations between the original ERV and nonrepetitive compartment are plotted as dots for reference, with 95% confidence interval.

We hypothesized that the CpG C>G mutational signature could result from the high and variable rates of CpG hydroxymethylation (hmC) of ERVs, which has been recently shown to increase rates of C>G mutations (Supek et al. 2014). To test this hypothesis, we compared the mutation spectra of ERVs with versus without evidence of hmC CpG, based on hmC-specific sequencing of human embryonic stem cells (Yu et al. 2012) (Table S2). ERVs with hmC showed a significant enrichment for CpG C>G mutations compared to ERVs without hmC in all six species, supporting the hypothesis of hmC-related mutagenesis in ERVs (Figure 4B, Table S6). We assessed the robustness of the CpG C>G mutational signature to differences in mapping quality, compartment size, and species-specific nucleotide content, finding that the CpG C>G mutational signature was robust to all quality control tests (Figure 4C, Figures S16-17).

### Conserved mutational signatures are associated with functional genomic elements in distantly related mammalian species

After observing that compartment-associated mutational signatures appear to be conserved among great ape species, we hypothesized that such conservation might extend to even more distantly related species and tested this hypothesis using annotations of epigenetic function that were generated by the ENCODE project for humans and mice. Specifically, we split the human and mouse genomes into six different functional compartments based on chromHMM annotations generated from epigenetic assays run in ESCs and mESCs, respectively (Ernst & Kellis 2012; Pintacuda et al. 2017) (Table S7). These chromHMM-defined compartments do not necessarily correlate with the various compartments we analyzed across great apes. We posited more generally that mutational signatures associated with specific compartments that experience specific molecular interactions (DNA binding proteins, histone modifications, chromatin state, e.g.) are relatively stable among similar genomic regions in different species. Across these compartments, we calculated normalized mutation spectra using publicly available whole-genome polymorphism data from 2504 diverse humans and 67 wild-caught *Mus musculus* and *Mus spretus* individuals (The 1000 Genomes Project Consortium 2015; Harr et al. 2016). Ancestral states for human polymorphisms were determined with regards to an inferred reconstructed genome representing the most recent common ancestor of humans; ancestral states for mouse were polarized based on an aligned rat reference genome (rn6) (The 1000 Genomes Project Consortium 2015). Polymorphisms were subject to filters similar to those applied to the great ape data: sites that either failed quality filters, were missing from >20% of haplotypes, or had a minor allele frequency of 1/2N (e.g. singletons) were excluded from analysis. Sites fixed in the sampled haplotypes were also excluded. We employed a similar randomization strategy to avoid structure due to shared variation, but summed together the mutation spectra of humans from the same subpopulation and mice from the same sampling site and subspecies to avoid sparseness (n=26, 9 respectively; see Methods).

Two separate matrices containing mutation spectrum data from all compartments in each respective species were decomposed using PCA, and we observed several commonalities between human and mouse in the spatial arrangement of the chromHMM compartments within the span of the first two principal components (Figure 5). Unsurprisingly, mouse species separate to a greater degree than do human populations, but within a subspecies the relative positioning of spectra from different functional compartments mirrors that observed in humans, especially after a roughly 30-degree rotation of the PC axes. In humans, PC1 largely separates promoters from other compartments, and PC2 separates transcribed regions. Enhancers and insulators cluster together, indicating mutation spectra more similar relative to other compartments. These same features are true of the mouse PCA, except that the vector separating promoters from other compartments is intermediate between PC1 and PC2. Mutation spectra from the heterochromatin and repressed compartments appear to have somewhat swapped places in humans and mouse; this phenomenon could be due to annotation differences. Furthermore, the loadings of PC1 and PC2 are significantly correlated between the two separate PCAs (Pearson’s ρ = 0.486, 0.555, p = 5.14*10^−7^, 4.44*10^−9^, respectively) (Figure S18). As expected, CpG transition rates are weighted heavily along PC1, likely reflecting the unique methylation patterns that regulate promoter function, but other types of transitions and transversions also appear to have different rates among these functional compartments. Both C>A and A>T, the mutation types associated with late replication timing in great apes, appear negatively associated with PC2, a pattern that is consistent with the tendency of transcribed regions, which are positively displaced along PC2, to occur in early replicating regions. Although ChromHMM-defined compartments are not necessarily orthologous between human and mouse, their shared epigenetic markings and functions appear to be enough to cause a common set of sequence motifs to be particularly vulnerable to mutation.

**Figure 5:**
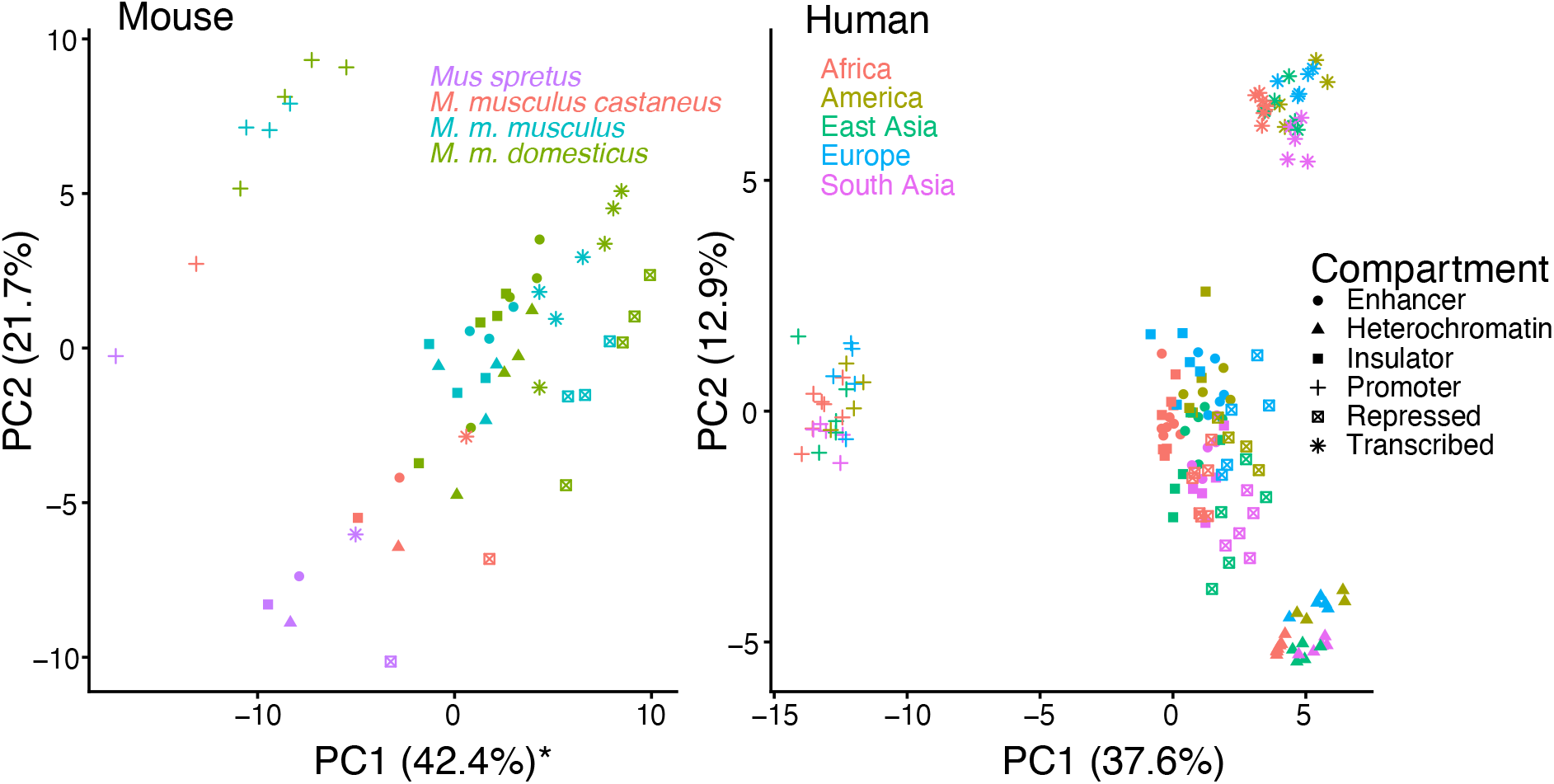
Structure of mutation spectrum variation is conserved between mouse and human. A-B. We ran PCA separately for mutation spectrum data from mouse (A) and human (b) individuals, whose genomes were split into compartments annotated by chromHMM (PC1 of mouse reversed for readability). The axes of greatest variance that separate clusters of compartment mutation spectra for both humans and mouse are highly similar. For both species, PC1 separates promoters from other compartments, while PC2 distinguishes transcribed regions. The loadings of PC1 and PC2 are significantly correlated between the two separate PCAs (rho = .486, 0.555, p = 5.14*10^−7^, 4.44*10^−9^, respectively) (Supplemental Figure S18).

## DISCUSSION

Despite considerable documented evidence of mutation spectrum variation across genomic space and phylogenetic time, little was previously known about the covariation of mutation spectra along spatial and temporal axes. Our results show that such covariation is negligible, at least in great apes: spatial and temporal mutation spectrum variation are largely orthogonal to one another. Replication timing, repeat content, and other functional categories have consistent mutational biases across all great ape species. At the same time, each species has a distinctive mutation spectrum bias that affects all functional compartments we have analyzed. There exist some exceptions to this general rule, most notably in the compartments of the genome that accumulate a maternal-age-related signature in humans that is attenuated in chimpanzees and gorillas and nearly nonexistent in orangutans. Nevertheless, our results show that mutation spectrum divergence between ape species is mostly driven by processes that act promiscuously across the genome. Determining how broadly this conclusion applies to species beyond great apes will be an exciting avenue for future work.

Our results provide evidence that mutation rates in late replicating regions are elevated due to a mechanism whose activity pattern over great ape evolution has been largely conserved. The distinct signatures associated with different great ape species demonstrate that the simplest possible “hominoid slowdown” model is likely not sufficient to explain all differences between great ape mutation rates. On one hand, some papers have suggested that increasing reproductive age is enough to explain differences in mutation rates that have been inferred by analyses of the branch lengths of great ape phylogenies. However, we see no evidence that differences between great ape species’ mutation spectra can be explained by varying the dosage of a single mutational signature associated with increased parental age, or even separate signatures associated with paternal and maternal ages of the kinds that have been inferred from human de novo mutation data.

To understand why species-specific mutational signatures are not obviously biased toward particular genomic regions, it is helpful to consider previous theoretical work on the dynamics of alleles that drive mutation rate evolution (Sturtevant 1937; Kimura 1967; Lynch 2008; Lynch et al. 2016). Although a genetic modifier of the mutation spectrum will not necessarily change the mutation rate, the most parsimonious scenario is for new mutations that alter the mutation spectrum to slightly increase the overall mutation rate by impairing the faithful replication of particular motifs. Kimura, Lynch and others have noted that selection against alleles that increase the mutation rate is driven by selection against the excess deleterious variation that such alleles beget. However, most new variants created by a *trans*-acting mutator will be on different chromosomes or distant parts of the same chromosome that will immediately recombine with other genetic backgrounds. The only deleterious mutations that are likely to cause selection against the mutator are mutations that happen to occur in a small window that maintains high linkage disequilibrium with the mutator locus. In contrast, a mutator that affects mostly neighboring DNA will tend to stay linked to more of the deleterious mutations it creates, making it more susceptible to purifying selection and loss from the population. A *trans* acting rate modifier that targets specific genomic regions might not experience strong linked selection itself, but the associated genomic targets might experience selective pressure to stop attracting targeted mutagenic activity.

Some species-specific signatures might be the footprints of environmental mutagens, but *trans*-acting genetic modifiers are more parsimonious explanations for signatures that affect larger clades of multiple species. The mutations we analyzed here are all segregating variants that originated long after modern ape species had become reproductively isolated, and environmental exposures are not likely to have respected phylogenetic boundaries for millions of years after the completion of ape speciation. Fixed differences between polymerases, DNA repair factors, and/or their regulatory elements are more likely to be responsible for differences in mutation spectra that respect phylogenetic structure and act consistently across the genome.

We have noted that all ape species exhibit some internal mutation spectrum substructure, with Western chimpanzees being the most distinctive subspecies. Western chimpanzees are different from other chimpanzee subspecies in several ways, including lower levels of bonobo gene flow, a higher load of transposable elements, and a stronger population bottleneck in their recent history. Both transposable elements and accelerated genetic drift may have hastened this lineage’s rate of mutation spectrum drift.

Although selection, drift, and demographic history certainly affect genome-wide genetic diversity and variation in diversity across the genome, these forces are not reasonable explanations for the patterns of mutation spectrum divergence we present in this manuscript. Biased gene conversion selects for mutations from A/T to G/C, but none of the mutational signatures that vary between compartments or species fit this simple profile. In coding regions, selection generally allows synonymous mutations to reach higher frequencies than nonsynonymous mutations, but coding regions comprise a nonexistent to negligible fraction of the various compartments analyzed here, and the universality of the genetic code prevents selection against nonsynonymous substitutions from generating any distinctive species-specific differences. It is generally not plausible to suppose that the same mutational signature would be selected for in a single lineage across all the noncoding genomic compartments we analyze here, as this would require mutations in many triplet contexts to have fitness effects that were somehow consistent across the whole genome.

Although identifying the causal fixed differences still represents a challenging unsolved problem, the insights from this paper will allow us to narrow the field of possible mutation spectrum modifiers to exclude ones that target only subsets of the genome. Focusing on ERVs in detail, we were able to use functional genomic annotations to link this compartment’s CG>GG mutational signature to hydroxymethylation of CpG sites. Examination of a broader set of functional genomic data may facilitate the interpretation of other localized signatures and bring us closer to understanding their causality.

Previous work estimated that 80% of spatial mutation rate variation could be explained by letting mutation rates depend on an extended 7-mer sequence context (Aggarwala & Voight 2016; Carlson, Locke, et al. 2018). Since 7-mer composition differs between genomic compartments, these extended sequence context models likely derive some of their predictive power from the effects of *cis*-acting mutational modifiers. However, we have shown that compartment annotations provide extra information about mutability above and beyond what we can tell from extended sequence context alone, at least in the case of the ERV hydroxymethylation signature. An important avenue for future work will be to examine the converse possibility and determine how much of the dependence of mutability on extended sequence context can be explained by genomic compartmentalization.

A small proportion of the genome is expected to vary in mutation rate between individuals due to the presence of rtQTLs, but our results suggest that the coarse shape of the replication timing landscape is stable across the great ape clade (Koren et al. 2012). The genomic distribution of the replication timing signature could even be leveraged to estimate the extent of TAD variation within and between species. More generally, if mutational signatures of chromatin states and various epigenetic modifications prove stable and interpretable over large phylogenetic clades, they represent a valuable source of information about the evolution of chromatin structure and function in non-model organisms where only genome sequences are available.

## METHODS

### SNV filtering

We ascertained mutation spectra from a set of high-coverage great ape SNVs that were previously called and filtered by Prado-Martinez et al. (Prado-Martinez et al. 2013). For each species and compartment, we collated the set of biallelic SNVs falling within the genomic segments that comprise that compartment. Ancestral states were assigned using a parsimony approach. Briefly, a biallelic site segregating within a genus (*Homo*, *Pan*, *Gorilla*, or *Pongo*) was polarized to the allele fixed in all other genera. Sites segregating in multiple genera, sites with multiple fixed alleles, and sites with more than two alleles in a single genus were excluded due to their inconsistency with the assumptions of no balancing selection and only one mutation event per site. Singletons were also excluded due to their higher likelihood of sequencing error. We used the inferred ancestral base to classify 3’ and 5’ neighboring nucleotides. For 3-mer mutational analyses, we excluded SNVs whose 3’ and 5’ neighboring nucleotides were an ‘N’ in the hg18 reference and SNVs demonstrating evidence of recurrent mutation in the great ape lineage; we expanded this filter to include the three 3’ and 5’ neighboring nucleotides for 7-mer mutational analyses. Finally, we removed SNVs out of Hardy-Weinberg Equilibrium with excess heterozygosity (using an exact test, *p*<0.05). Excess heterozygosity at a locus could indicate a cryptic segmental duplication with a single, fixed mutation in a copy. We also excluded SNVs with ≥0.5 derived allele frequency to avoid mutational classes with an elevated risk of ancestral state misidentification.

### Computing the mutation spectra of individuals and species

The PCA analyses in this paper require the computation of mutation spectra from individual genomes, whereas the complementary heat map analyses involve calculating aggregate mutation spectra from larger samples. Each analysis employs the filtering system described above and ultimately involves counting the number of filtered derived alleles, classified by 3-mer context. However, slightly different calculation details are involved in the two cases.

The aggregate mutation spectrum of a species *S* is obtained from a set of counts *C*(*m_1_,S*),…,*C*(*m_96_,S*) where *m_1_*,…,*m_96_* are the 96 3-mer mutation type categories AAA>C, …, TCT>T. The count *C*(*m_1_,S*) is the total number of SNVs segregating in species *S* that fall into the mutational equivalence class *m_1_*. To compare spectra across samples with different amounts of variation, these mutation type counts are normalized to obtain a 96-dimensional histogram with frequency categories summing to 1.

A mutation spectrum can be similarly calculated from a particular individual *I* as the distribution of 3-mer mutation types across the derived alleles present in *I*’s genome. Homozygous derived alleles are given twice the weight of heterozygous alleles such that the spectrum is the average of the spectra one would compute from the two phased haplotypes making up *I*’s diploid genome.

When individual mutation spectra are computed in this way, two types of derived alleles can contribute to spectrum covariance between individuals *I* and *J*. The first type are pairs of derived alleles that occur at separate loci in *I* and *J* but belong to the same mutation equivalence class. The second type are derived alleles inherited by both *I* and *J* from a common ancestor. To maximize our power to detect mutation spectrum evolution and distinguish it from shared genetic drift, we devised a randomization strategy to eliminate the second source of signal while preserving the first.

This randomization strategy involves computing the mutation spectrum of individual *I* from only a subset of the derived alleles present in *I*’s genome. If one copy of a particular derived allele is present in *I*’s genome and has frequency *k/2N* in the GAGP panel, the allele will be counted toward *I*’s mutation spectrum with probability 1/*k*. Conversely, this derived allele will be counted toward the mutation spectrum of exactly one ape haplotype that carries it, with the identity of that haplotype chosen uniformly at random.

### Comparing mutation spectra across genomic compartments

Comparing mutation spectra between regions of the genome required accounting for differences in compartment size and nucleotide content. Larger compartments naturally had more mutations than smaller ones; it was therefore necessary to compare mutation fractions rather than raw counts. Furthermore, differences in nucleotide content between compartments could bias our comparison and calculation of local mutation rates. For example, a particular compartment could have a relatively high count of AAA>ACA SNVs, but this high count might be caused by the compartment having many occurrences of the triplet AAA, and therefore more opportunities for an AAA>ACA to occur. Thus, we rescaled the number of mutations for each compartment by the nucleotide content of the NRNC compartment before calculating fractions. To calculate the rescaled rate *r*(*m*) of mutation *m_i_*: {*m_1_*, …, *m_96_*} corresponding to triplet *t_t_*: {*t_1_*, …, *t_32_*} and compartment *C*:

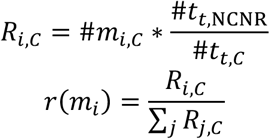

We calculated triplet content of each compartment by sliding a 3bp window, 1bp at a time, across each compartment. Edges of compartment segments and triplets with N’s were excluded.

Several statistical analyses comparing two different mutation spectra required count data rather than frequency data (e.g., Chi-square tests). We devised a slightly different rescaling strategy for these counts to avoid artificially inflating the mutation counts. To calculate the rescaled count of mutation *m_i_*: {*m_1_*, …, *m_96_*} corresponding to triplet *t_t_*: {*t_1_*, …, *t_32_*} and compartment *C_1_* in preparation for comparison to compartment *C_2,_* we scaled down the raw count of mutation *m* in the compartment where *m* is more abundant, rather than scaling up the count of *m* in the compartment where it is more abundant:

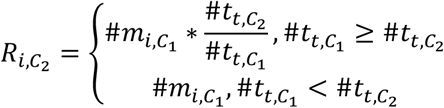

The same rescaling is used for compartment *C_2_*, switching the subscripts accordingly.

### Statistical analyses

We generated plots and performed statistical analyses in R (version 3.1.0) using scripts available at https://github.com/harrispopgen/gagp_mut_evol.

We ran **PCAs** on matrices (*k* × *C* rows by 96 columns, for *k* individuals and *C* compartments) of rescaled 3-mer mutation rates calculated for each individual and each compartment using the *prcomp* method. Some PCAs were run on individuals from all species; others were only run on individuals from a single species. The matrices were centered and scaled, as is standard for *prcomp*. The **PCA loading heatmaps** display the weights associated with each of the 96 3-mer mutation types for a given PC. The **Euclidean distance ridge plots** were similarly generated with individual mutation spectrum data that required no rescaling since the analysis considered only a single compartment. We plotted the distribution of Euclidean distances based on a 2 × 96 matrix comparing the mutation counts between two different individuals of either a single or two different species. For comparisons within species, we ran *k* choose 2 tests (*k* being the number of individuals for a given species); for comparisons between two species, we ran *k* × *l* tests (*k* and *l* being the numbers of individuals in both species respectively).

The **log-odds heatmaps** were generated to display the relative enrichment or depletion of specific mutation types when comparing two compartments directly to each other. For a given species, we plotted the log transform of the ratio between the rescaled mutation rates of two compartments.

The **7-mer content-corrected heatmap** required 7-mer mutation and nucleotide content from various compartments, which were generated using the 3-mer mutation and nucleotide methods and equally expanding context on the 5’ and 3’ side of the central base. Each original 3-mer mutation *m_3,k_*: {AAA>ACA, AAA>AGA … TCT>TTT} is a collapsed equivalence class of 256 unique 7-mer mutations *m_7,i,x_*: {AAAAAAA>AAACAAA, AAAAAAC>AAACAAC, … TTAAATT>TTACATT}. To explicitly re-weight the counts of each 3-mer mutation *m_3,i,C_* in compartment *C* using the ratio of 7-mer content in *C s_s,C_*: {AAAAAAA, AAAAAAC, … TTTCTTT} to that of compartment *C’*:

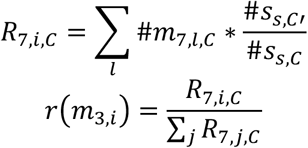

We used this method to rescale 3-mer mutations from the ERV compartment to match the 7-mer content from the nonrepetitive heterochromatin compartment; the heatmap in Figure S17 presents the log ratio of the rescaled ERV mutation and the non-rescaled nonrepetitive heterochromatin mutation spectra.

We ran **correlation analyses** to quantify the similarities between the mutation spectrum heatmaps between species. Each mutation spectrum heatmap comprised the log ratio between each of the 96 mutation types between two compartments for a single species. To calculate the similarity between heatmaps for two different species, we ran a Pearson correlation test on the paired vectors of log odds.

### Compartments

We defined compartments based on published annotations of genomic features. Each compartment was a list of genomic segments in a bed file format. The following is a list of the compartments used in our analyses and a short description of how we generated them.

**NCNR**: (non-conserved, non-repetitive) the entire hg18 genome, excluding repetitive elements defined by repeatMasker, conserved regions in primates based on the phastCons 44-way multi-species alignment, CpG islands from the UCSC genome browser, and coding exons from refGene.
**ERVs**: all ERVs in repeatMasker run on hg18, excluding those classified specifically as mammalian long-terminal repeats (MaLRs). MaLRs are believed to be largely inactive in great apes, unlike ERVs which are still active in several species.
**LINEs**: all LINEs in repeatMasker run on hg18
**Heterochromatin**: the intersection of the heterochromatin domain called in the hg18 chromHMM run on 9 different cell types from ENCODE (Gm12878, H1 HESCs, HepG, Hmec, Hsmm, Huvec, K562, Nhek, and Nhlf), minus repetitive elements defined in repeatMasker.
**Early/late replicating regions**: the genomic quartiles that replicate earliest and latest during S phase were ascertained using replication timing data from Koren et al., 2012 (Koren et al. 2012). In that manuscript, the fine-scale replication timing of regions in the genome was determined by sequencing human lymphoblastoid cell lines at S1/G phase; read depth over a region in the genome corresponded to its average relative replication timing. The read depths were measured at specific genomic positions. We calculated average replication timing for non-overlapping 20kb window that included at least one measurement of replication timing and calculated replication timing quartiles. The earliest and latest quartiles were used as compartments. **Early/late replicating, repetitive/non-repetitive regions** were the subsets of the replication timing compartments that overlapped with or excluded repeatMasker-annotated repeats, respectively.
**Human maternal mutation hotspots**: defined in Jonsson et al., 2017 as regions whose *de novo* mutation rate strongly associates with maternal age, lifted over to hg18 (Jónsson et al. 2017).
**ERVs ± 5hmC**: ERV compartment, split into segments that had or lacked evidence of ≥ 1 5hmC site, based on Tet-assisted bisulfite sequencing of human ESCs (Yu et al. 2012).

### Quality control analyses

We ran a number of analyses to test the robustness of our methods and findings.

#### Determining how GC-biased gene conversion contributes to the separation of species and compartments in PCA plots

GC-biased gene conversion (gBGC) is the nonreciprocal copying of short DNA tracts between homologous chromosomes during meiosis, which at heterozygous sites tends to retain G/C (strong, S) alleles more often than A/T (weak, W) alleles (Eyre-Walker 1993, 1999; Holmquist 1992). This process causes G/C derived alleles to have higher substitution rates and frequencies than A/T alleles. If the strength of gBGC were not conserved across the great ape phylogeny, we might expect species’ and compartments’ mutation spectra to separate along an axis defined by the ratio of W-to-S and S-to-W mutation types, with species and compartments that experience the most gene conversion having the highest proportions of S-to-W variants. Such a gradient would be expected to dissipate, however, if we constructed mutation spectra using a higher proportion of rare variants, which are younger than common variants and have had less time to be influenced by the effects of gBGC. To this end, we tabulate mutation spectra in the following way: each SNV present in *k* haplotypes is counted toward the mutation spectrum for only one randomly selected haplotype; this process is described above (“computing the mutation spectra of individuals and species”) and hereafter referred to as “randomization.” We observe that individuals cluster by species in a PCA run on NCNR mutation spectra whether we employ randomization or count each SNV toward every haplotype on which the derived allele appears. Furthermore, the principle component loadings are not consistent with the expected signature of gBGC. Many of the mutation types that have different mutation fractions in different ape species are W-to-W or S-to-S mutation types that are expected to be unaffected by gBGC (Figure S2). Even if we compute mutation spectra entirely from rare doubleton variants, we observe similar species separation to what we observe using more common variants, despite these rare variants being more weakly affected by gBGC.

The rates of gBGC covary across the genome with recombination rate. The locations of recombination and gBGC hotspots change rapidly and are often species-specific due to evolution of the gene *PRDM9*. Thus, comparing mutation spectra between species in locations where gBGC rates are most divergent should (A) indicate an upper bound on the effects of gBGC rate evolution on our mutation spectrum analyses and (B) test our capacity to minimize its effects through the randomization method. We therefore examined the mutation spectra of species-specific recombination hotspots with and without our random sampling method (Figure S1). These analyses showed that, without random sampling, differences in mutation spectra within recombination hotspots are in fact dominated by the *cis*-acting effects of GC-biased gene conversion (gBGC). Figure S1A shows a PCA of individual mutation spectra from the NCNR compartment and two compartments containing the genomic regions whose recombination rates fall within the upper and lower genome-wide deciles in humans (Kong et al., 2010); this spectrum is not thinned by randomization. We can clearly see the mutation spectra cluster by species, but we can also see high-recombination-rate and low-recombination rate compartments separate along PC2, especially in humans where the ascertained recombination hotspots are active. PC2’s loadings are dominated by A>G and A>C mutations, which are both W>S mutations and likely indicate a human-specific enrichment for gBGC. In contrast, the separation between high recombination and NCNR compartments is mostly attenuated in Figure S1B with the use of the randomization method.

#### Comparing SNV to de novo mutation spectra

Several papers have recently reported homogeneity among the de novo mutation (DNM) spectra of great apes. Our analyses, however, demonstrate that mutation spectra generated from SNVs vary significantly between species. We therefore compare our SNV to DNM spectra from Besenbacher et al. (2019) to assess the difference in results and find that DNM spectra are underpowered to detect the species signatures we find in SNV spectra.

We test the likelihood that the distribution of the observed DNM counts in an individual is pulled from the SNV spectrum of a given species *s*. We calculate the expected probabilities *p* in [*p_A>C,s_, p_A>G,s_, … p_C>T,s_*], given the number of SNVs segregating in species *s* in the GAGP of mutation type *m_i,s_* in [*m_A>C_*, *m_A>G_*, … *m_C>T_*] by the following equation:

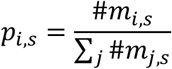

We then calculate the log likelihood that the spectrum of DNMs for individual *x* (*m_i,x_*) assuming a multinomial distribution. This likelihood represents how well the DNM spectrum for an individual fits the SNV spectrum of a given species.

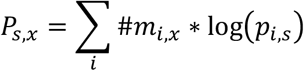

To determine whether a SNV spectrum fit a DNM spectrum significantly better than others, we calculated the significance of the differences in likelihoods for a given individual when compared to chimpanzees, orangutans, and gorillas (*x* : [*C, O, G*]). We simply calculated the fold range in fit as:

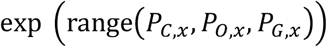

A “significance” threshold was set at a fold range of 20x, to replicate a standard *α* value of 0.05. The table below shows that none of these values approach 20 (Table S1). We conclude that the DNM spectra are too sparse to demonstrate clear species differences.

#### Testing the robustness of PCA clustering to species representation

We tested the robustness of the clustering of individuals by species in the NRNC PCA to differences in number of individuals sequenced per species. We down-sampled the number of individuals per species to match those of humans (*n* = 9), excluding the two orangutan species who were grouped as a single super-species group for this analysis. The clustering patterns in the PCA remained.

We recreated the several plots from Figs. 1 and 2 using a rarer subset of variants and found no qualitative differences when compared to the original figures (Figure S6A). To avoid the potential quality issues related to relying on singletons, we used only doubletons (DAF = 2/2*N*, for *N* individuals sequenced from each species) in this analysis. The PCA of the individual mutation spectra across all species in the NCNR compartment alone still demonstrates the clustering of individuals by species (Figure S6B). The distribution of Euclidean distances between the NCNR mutation spectra of individuals within and between species still demonstrates the correlation of mutation spectrum distance with divergence time (Figure S6B). The PCA of individual mutation spectra from the NCNR, early replication timing, and late replication timing compartments still demonstrates similar trends as observed in figure 2D (Figure S6C). The first PC, representing the greatest axis of variance among all spectra, separates spectra according to phylogeny and represents species signatures; the second PC separates compartments by their replication timing. The loadings for PC2 are enriched for C>A and A>T mutation types, corresponding with the known late replication timing signature (Figure S6D). We attribute the spread observed in chimpanzees in the replication timing doubleton PCA below to be largely an effect of noise imposed by down-sampling.

#### Testing the CG>GG signature in ERVs

We determined the enrichment for CG>GG mutations in ERVs compared to nonrepetitive heterochromatin was unaffected by the differences in mapping quality between the two compartments, noting that mutations in repetitive regions are more difficult to call confidently. To determine the potential confounding effect of mapping quality on the CG>GG signature, we compared the distribution of the mapping quality value of the variants in the ERV and nonrepetitive heterochromatin compartments (MQ field in the GAGP vcf). Density distributions of the mapping qualities for the two compartments were highly overlapping within each species (Figure S20).

We also determined that the CG>GG signature was unaffected by the different ‘shapes’ of the ERV and nonrepetitive heterochromatin compartments, i.e. the distribution of segment lengths and overall size of compartment. For each ERV compartment segment for a given chromosome, we reassigned the coordinates randomly within the nonrepetitive heterochromatin compartment, preserving segment length. In the event a randomized compartment was chosen to overlap one or more “N” bases, a new compartment was resampled. We did not filter for overlapping randomized segments, but assumed that, given that the ERV compartment was a fraction of the size of the heterochromatin compartment (127 Mb and 430 Mb respectively), collisions would be unlikely enough to that their sparsity would not bias our findings. This randomization process to create ‘ERV-like’ compartments was bootstrapped 100 times. We then calculated nucleotide content and mutation spectra for each of the 100 bootstrapped compartments. We generated the log-odds of the CG>GG mutation type between the bootstrapped compartments and nonrepetitive heterochromatin compartment (each normalized to each other, using the Chi-square normalization method as described above), then compared those values to the same statistic of the original two compartments (Figure 4C) for all species. The observed CG>GG enrichment in the ERV compartment lies outside of the null distribution in all species except Bornean orangutans, for which the confidence interval of the observed log-odds overlaps the null distribution. Given that the observed CG>GG fraction still falls outside of the null distribution, we conclude that Bornean orangutans show the same trend as seen in other species, but that trend is not significant on its own. This lower enrichment in Bornean orangutans may be a result of their low sample size (2N = 10) and the paucity of observed segregating variation (65% that of Sumatran orangutans, 30% that of gorillas, e.g.). The enrichment for CG>GG mutations comparing ERVs to nonrepetitive heterochromatin is significantly stronger than the enrichment in any of the bootstrapped, ‘ERV-like’ compartments (Table S6).

We tested the effect of species-specific nucleotide content on our rescaling method. The GAGP data are aligned to hg18; therefore, mutations in genomic regions in a non-human species that do not exist or do not map well in humans (e.g., new repetitive elements) were absent from our analyses. Our mutation rescaling method, however, relies on compartment nucleotide content determined from the hg18 reference, therefore including regions specific to humans and absent in other species. We compared the log-odds of each mutation type between the ERV and nonrepetitive heterochromatin compartment in all six species with those calculated by rescaling mutation counts using the nucleotide content of the segments of a given compartment that successfully lifted over to each respective species (lifted from hg18 to gorGor4, panPan1, panTro2, and ponAbe2 for gorilla, bonobo, chimpanzee, and both orangutans, respectively, using default liftOver settings). The log-odds values within species are highly significantly correlated (all *ρ* ≥ 0.95, Figure S19).

## DATA AVAILABILITY

No new data were generated or analyzed in support of this research. The processed mutation spectra and analysis pipelines are available in a public GitHub repository at https://github.com/harrispopgen/gagp_mut_evol.

## ACKNOWLEDGEMENTS

This work was supported by the National Institute of General Medical Science at the National Institutes of Health (T-32 GM081062 to M.E.G., 1R35GM133428-01 to K.H.); the Burroughs Wellcome Fund (a Career Award at the Scientific Interface, to K.H.); the Pew Charitable Trusts (Biomedical Scholarship, to K.H.), the Searle Scholars Program (Career Award, to K.H), and the Alfred P. Sloan Foundation (Research Fellowship, to K.H). We thank Evan Eichler, Phil Green, Sharon Browning, and members of the Harris lab for helpful discussions. We also thank Noah Snyder-Mackler and Aylwyn Scally for manuscript comments and Shwetha Murali for technical assistance.

