## Supplemental for "Mutational signatures of replication timing and epigenetic modification persist through the global divergence of mutation spectra across the great ape phylogeny"

SUPPLEMENTARY TABLES

| Genus | Species | Subspecies | Common name | Population | N |
|---|---|---|---|---|---|
| Homo | sapiens | - | Human | African | 3 |
| Homo | sapiens | - | Human | Non-African | 6 |
| Pan | troglodytes | ellioti | Nigeria-Cameroon chimpanzee | - | 10 |
| Pan | troglodytes | schweinfurthii | Eastern chimpanzee | - | 6 |
| Pan | troglodytes | troglodytes | Central chimpanzee | - | 4 |
| Pan | troglodytes | verus | Western chimpanzee | - | 5 |
| Pan | paniscus | - | Bonobo | - | 13 |
| Gorilla | beringei | graueri | Eastern lowland gorilla | - | 3 |
| Gorilla | gorilla | diehli | Cross river gorilla | - | 1 |
| Gorilla | gorilla | gorilla | Western lowland gorilla | - | 27 |
| Pongo | abelii | - | Sumatran orangutan | - | 5 |
| Pongo | pygmaeus | - | Bornean orangutan | - | 5 |

Table S1: Distribution of analyzed individuals in the GAGP by genus, species, subspecies, and population. Individuals were sequenced to a mean 25-fold coverage (Prado-Martinez et al., 2013).

| Compartment | bp (hg18) |
| --- | --- |
| NCNR | 1230819358 |
| Early replication | 683958817 |
| Late replication | 610749359 |
| ERV | 126631021 |
| Heterochromatin | 429839988 |
| LINE | 552512327 |
| Early replication, non-repetitive | 349059126 |
| Early replication, repetitive | 330655407 |
| Late replication, non-repetitive | 311710121 |
| Late replication, repetitive | 296248298 |
| Human recombination coldspots | 254546577 |
| Human recombination hotspots | 254370558 |
| ERVs, hmC+ | 16318060 |
| ERVs, hmC- | 110312961 |
| Human maternal hotspots | 234167393 |
| CpG islands | 29082042 |

Table S2: Sizes of compartments examined in great apes.

| Individual | Species | Max. diff. in likelihood |
|---|---|---|
| Aris | Orangutan | 2.82 |
| Carl | Chimpanzee | 1.39 |
| Dennis | Chimpanzee | 2.61 |
| Dylan | Chimpanzee | 1.46 |
| Efata | Gorilla | 1.10 |
| Marlies | Chimpanzee | 1.48 |
| Marlon | Chimpanzee | 1.31 |
| Mutasi | Gorilla | 1.09 |
| Pat | Chimpanzee | 1.66 |
| Ruud | Chimpanzee | 3.09 |

Table S3: Fold-differences in goodness-of-fit for the de novo mutation (DNM) spectrum of each individual from Besenbacher et al. (2018) with single nucleotide variant (SNV) spectra from orangutans, gorillas, and chimpanzees. The mutations in DNM spectra are too sparse to confidently distinguish between the goodness-of-fit to any particular species' SNV spectrum.

| Species 1 | Species 2 | $\rho$ | P-value |
|---|---|---|---|
| Human | Chimpanzee | 0.82 | 5.31E-25 |
| Human | Bonobo | 0.81 | 1.13E-23 |
| Human | Gorilla | 0.77 | 5.11E-20 |
| Human | Sumatran orangutan | 0.89 | 5.19E-33 |
| Human | Bornean orangutan | 0.84 | 1.59E-26 |
| Chimpanzee | Bonobo | 0.94 | 9.27E-45 |
| Chimpanzee | Gorilla | 0.94 | 1.80E-46 |
| Chimpanzee | Sumatran orangutan | 0.91 | 1.24E-37 |
| Chimpanzee | Bornean orangutan | 0.85 | 4.69E-28 |
| Bonobo | Gorilla | 0.93 | 5.97E-42 |
| Bonobo | Sumatran orangutan | 0.93 | 1.83E-41 |
| Bonobo | Bornean orangutan | 0.91 | 1.12E-37 |
| Gorilla | Sumatran orangutan | 0.87 | 5.12E-31 |
| Gorilla | Bornean orangutan | 0.86 | 9.76E-29 |
| Sumatran orangutan | Bornean orangutan | 0.95 | 1.54E-47 |

Table S4
A table of Pearson's correlation tests of the log ratios of mutation types comparting late to early replication timing compartments between each pair of species. P-values are uncorrected.

| Species 1 | Species 2 | $\rho$ | P-value |
|---|---|---|---|
| Human | Chimpanzee | 0.96 | 1.49E-16 |
| Human | Bonobo | 0.97 | 5.98E-17 |
| Human | Gorilla | 0.94 | 5.93E-14 |
| Human | Sumatran orangutan | 0.91 | 1.09E-11 |
| Human | Bornean orangutan | 0.93 | 3.86E-13 |
| Chimpanzee | Bonobo | 0.99 | 2.14E-22 |
| Chimpanzee | Gorilla | 0.98 | 9.75E-21 |
| Chimpanzee | Sumatran orangutan | 0.96 | 8.02E-16 |
| Chimpanzee | Bornean orangutan | 0.97 | 1.10E-16 |
| Bonobo | Gorilla | 0.98 | 7.64E-20 |
| Bonobo | Sumatran orangutan | 0.96 | 5.41E-16 |
| Bonobo | Bornean orangutan | 0.98 | 7.89E-19 |
| Gorilla | Sumatran orangutan | 0.96 | 1.59E-15 |
| Gorilla | Bornean orangutan | 0.96 | 2.78E-16 |
| Sumatran orangutan | Bornean orangutan | 0.98 | 2.29E-21 |

Table S5
PCAs run on individual mutation spectra from 8 different compartments are highly correlated among species, demonstrating uniform dosages of species mutational signatures. We calculated the midpoint of all individuals' mutation spectra for each compartment in PC1-3 and for each species. We then calculated the distance between the midpoints for each pair of compartments, computing a total of $\binom{8}{2} = 24$ distances for each species. We ran Pearson correlation tests between the paired vectors of distances between each combination of species, performing $\binom{6}{2} = 15$ tests; uncorrected P-values and estimates of $\rho$ are listed above.

| Species | P-value |
|---|---|
| Human | 1.90E-05 |
| Chimpanzee | 2.90E-12 |
| Bonobo | 2.03E-07 |
| Gorilla | 3.58E-09 |
| Sumatran orangutan | 0.00245204 |
| Bornean orangutan | 0.00014367 |

Table S6
Table of the P-values from a Chi-square test for different rates of CG>GG mutation types comparing the ERV hmC+ to hmC- compartments. P-values are uncorrected.

| Compartment | Size (human, hg19) | Size (mouse, mm10) |
|---|---|---|
| Promoter | 45669288 | 37074578 |
| Enhancer | 109737822 | 112146530 |
| Insulator | 21727890 | 18373998 |
| Transcribed | 106414354 | 148527127 |
| Repressed | 36676448 | 181894919 |
| Heterochromatin | 1845415220 | 146461481 |

Table S7: Sizes of chromHMM compartments in mouse and human in bp.

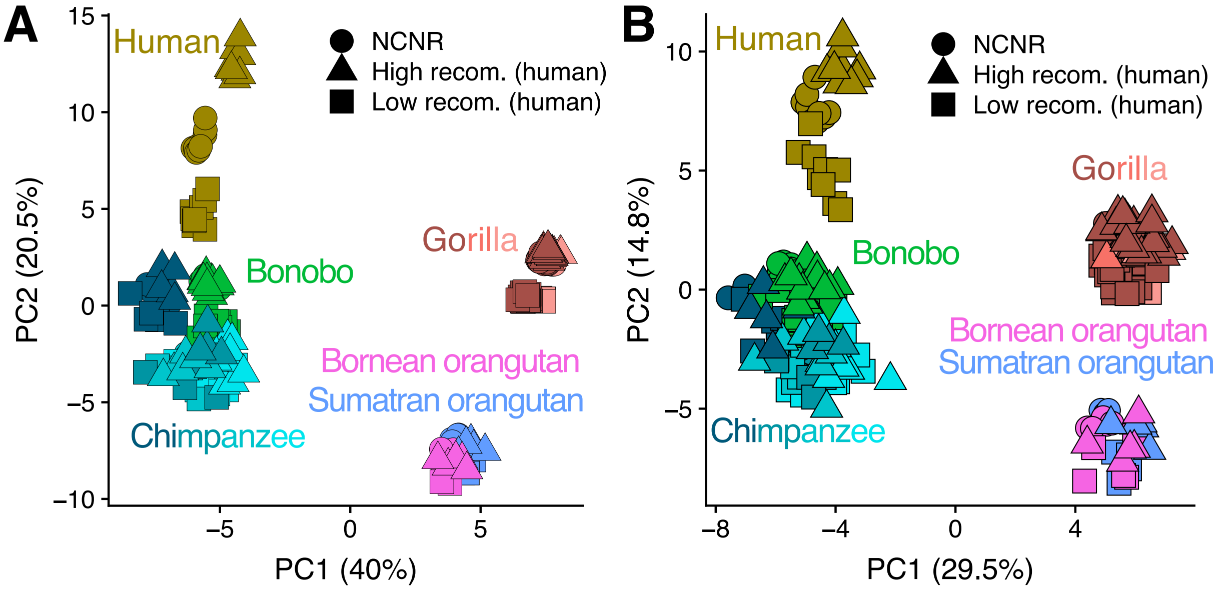


Figure S1: The effects of gBGC on mutation spectrum variation in recombination hotspots and coldspots is minimized by randomization method. We defined two compartments as those representing the top and bottom 10% of the genome ranked by local recombination rate in humans. The SNVs included in the spectra presented in (B) were randomly assigned to a single haplotype; this method was not applied in (A). The recombination hotspot mutation spectra in humans separate cleanly in (A); this separation decreases with divergence time from humans. These trends align with the rapid evolution of recombination hotspots, and therefore local rate of gBGC. The differentiation of recombination hotspot mutation spectra in (B) nearly disappears with the randomization method, thus demonstrating its minimizing effect on gBGC.

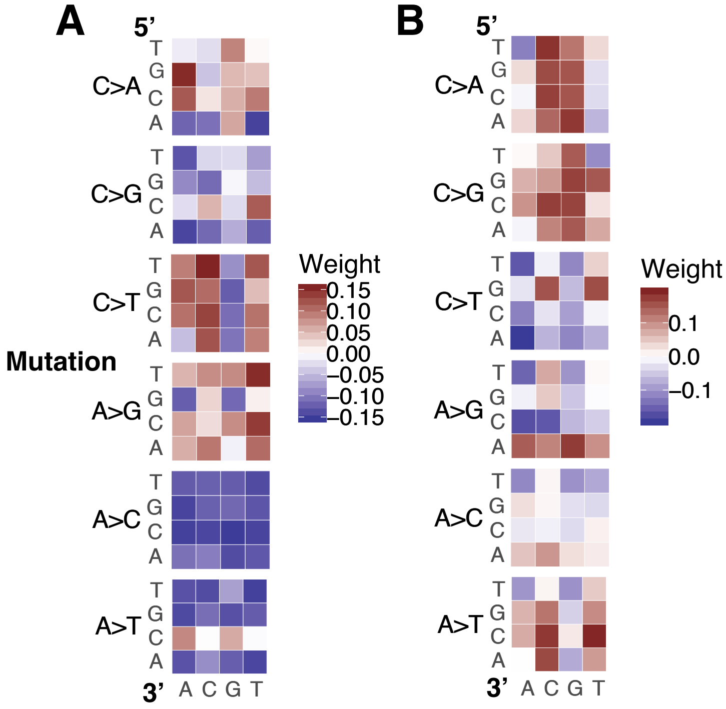


Figure S2: The loadings of PC1 (A) and PC2 (B) from PCA of NCNR spectra across all 88 individuals in the GAGP.

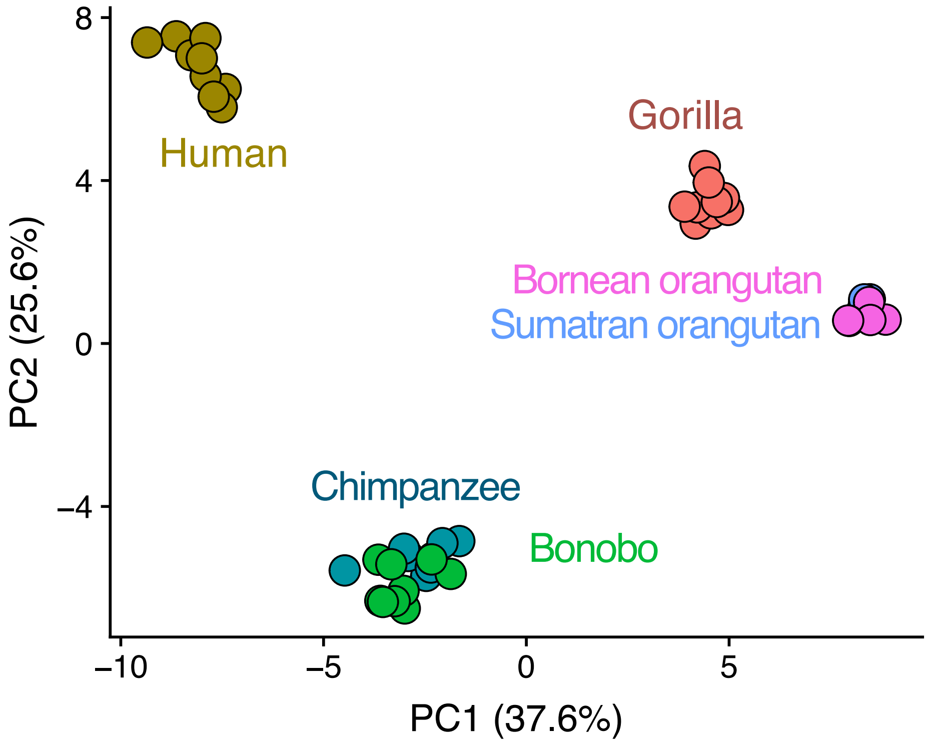


Figure S3
NCNR PCA clustering is robust to differences in species representation. We down-sampled each species (or both orangutan species, grouped together) to nine individuals and generated a PCA.

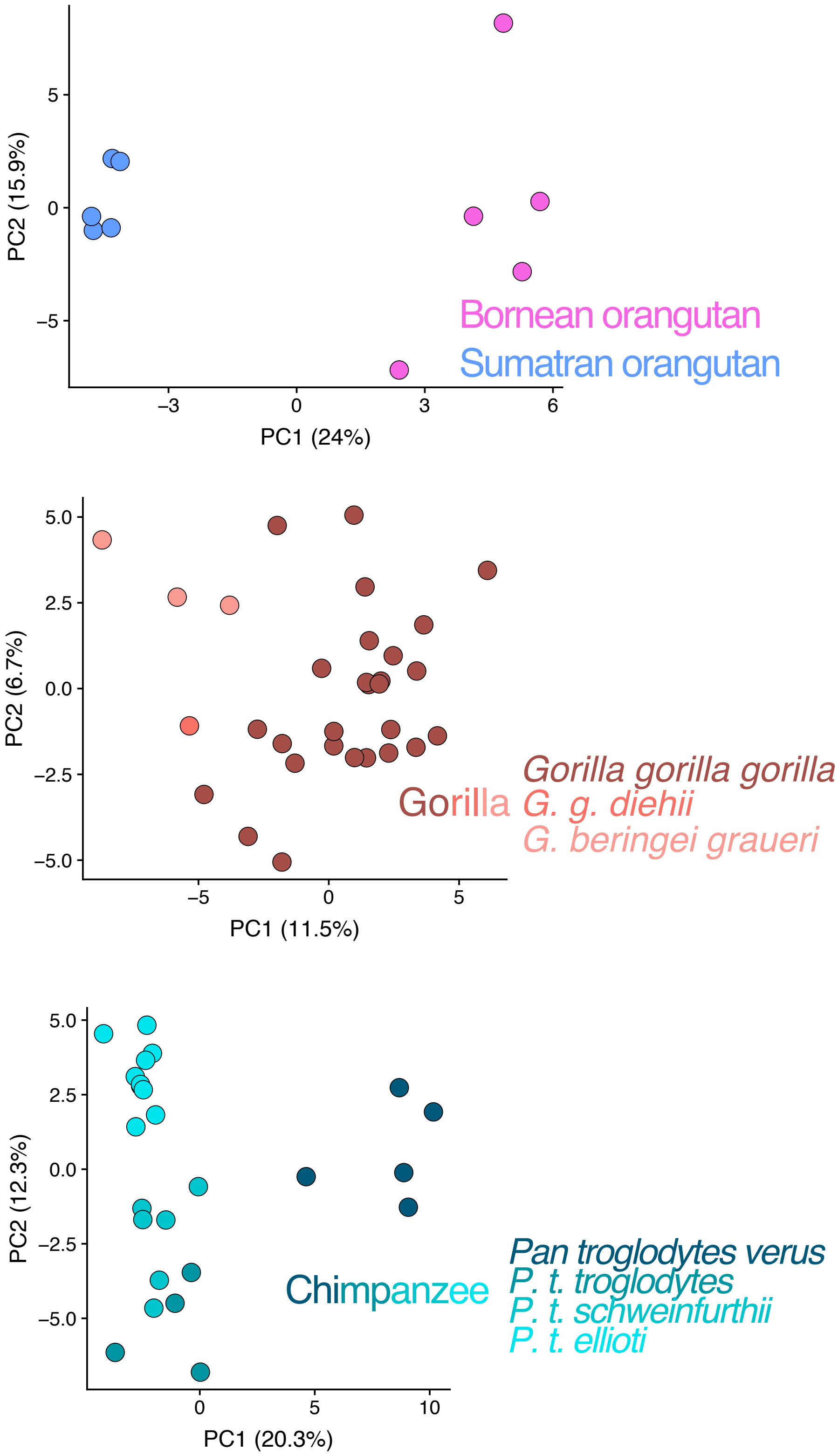


Figure S4
PCAs of the NCNR compartment for the orangutan clade, gorillas, and chimpanzees demonstrate finer-scale separation and clustering of individuals by subspecies.

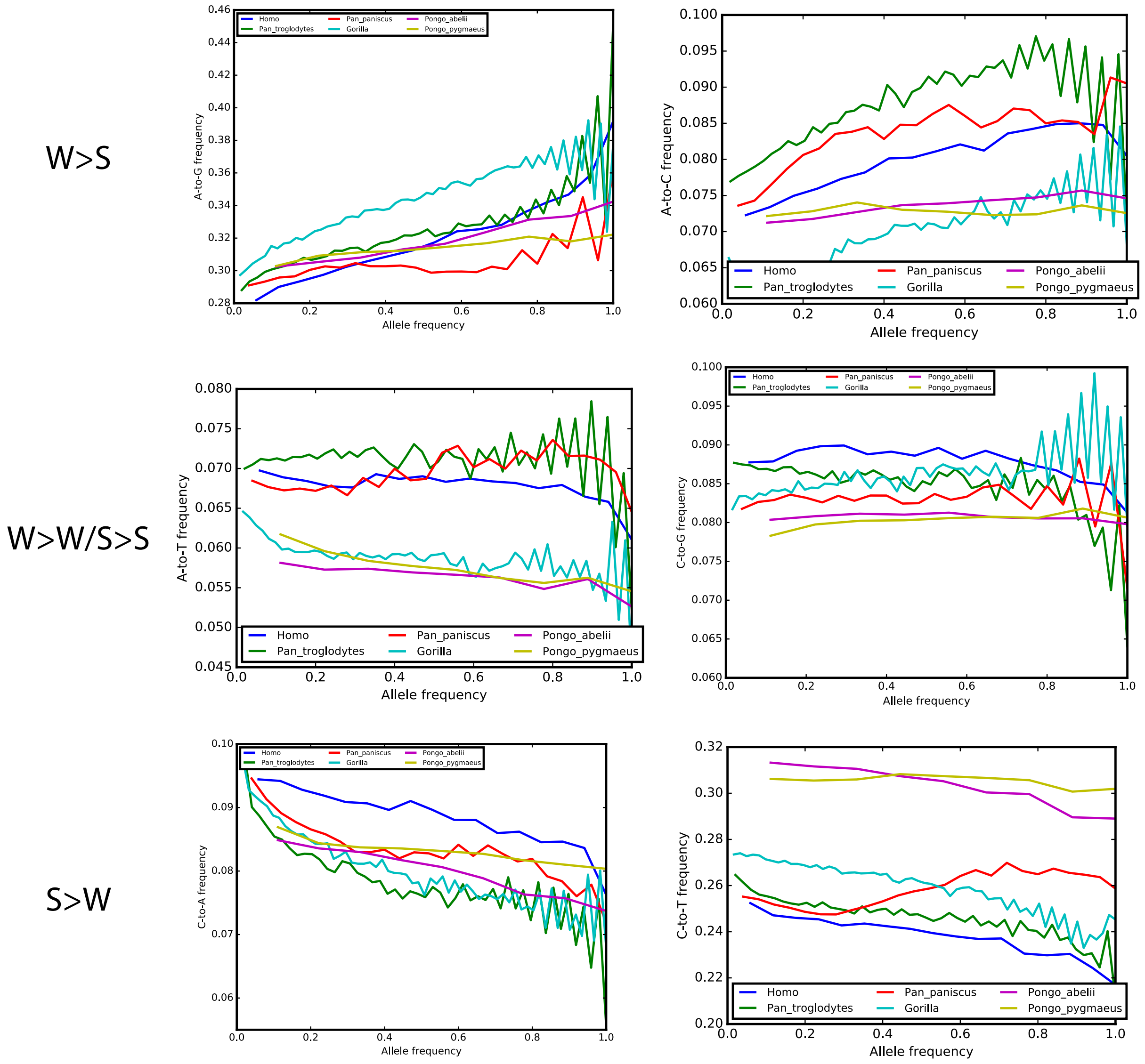


Figure S5
Effects of gBGC on allele frequency remain constant across great ape phylogeny. These graphs show the each species' composition of SNVs by different mutation types at each allele frequency. In general, high allele frequency SNVs show enrichment for W>S mutations and depletion for S>W mutations; this trend, in concordance with the effects of gBGC, is consistent across all species. Differences in the y intercept indicate differences in de novo mutation rates. Differences in the smoothness of the line, particularly in orangutan species, can be attributed to the low number of sampled haplotypes (2N = 10).

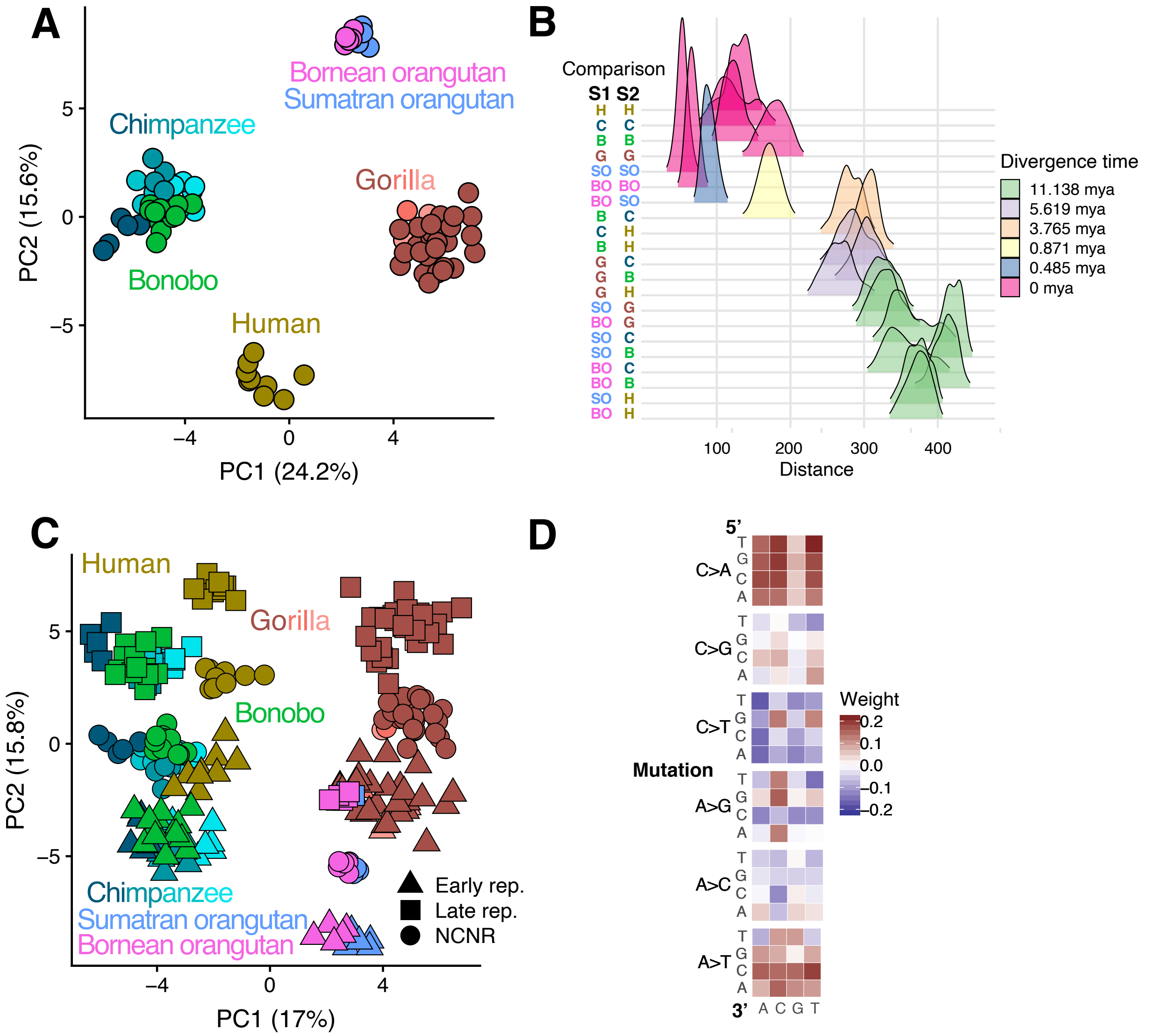


Figure S6: Rare variants demonstrate similar mutation spectrum trends to common variants

A. NCNR mutation spectra for 88 individuals in the GAGP cluster by species, demonstrating rapid, lineage specific evolution of mutation spectra. Different colors imply different species, and different shades of chimpanzees and gorillas are different subspecies.
B. Euclidean distances between mutation spectra of intra- and inter-specific pairs of individuals demonstrates an association of mutation spectrum distance with divergence time.
C. PCA of mutation spectra from NCNR, early replication, and late replication timing compartments for 88 individuals shows separation along orthogonal "phylogenetic" and "replication timing" axes.
D. The loadings of PC2, which roughly corresponds to the "replication timing" axis, shows an enrichment for C>A and A>T mutation types. Late replication timing regions are known to be enriched for this mutation signature; this recapitulates results from using rare and common variants together.

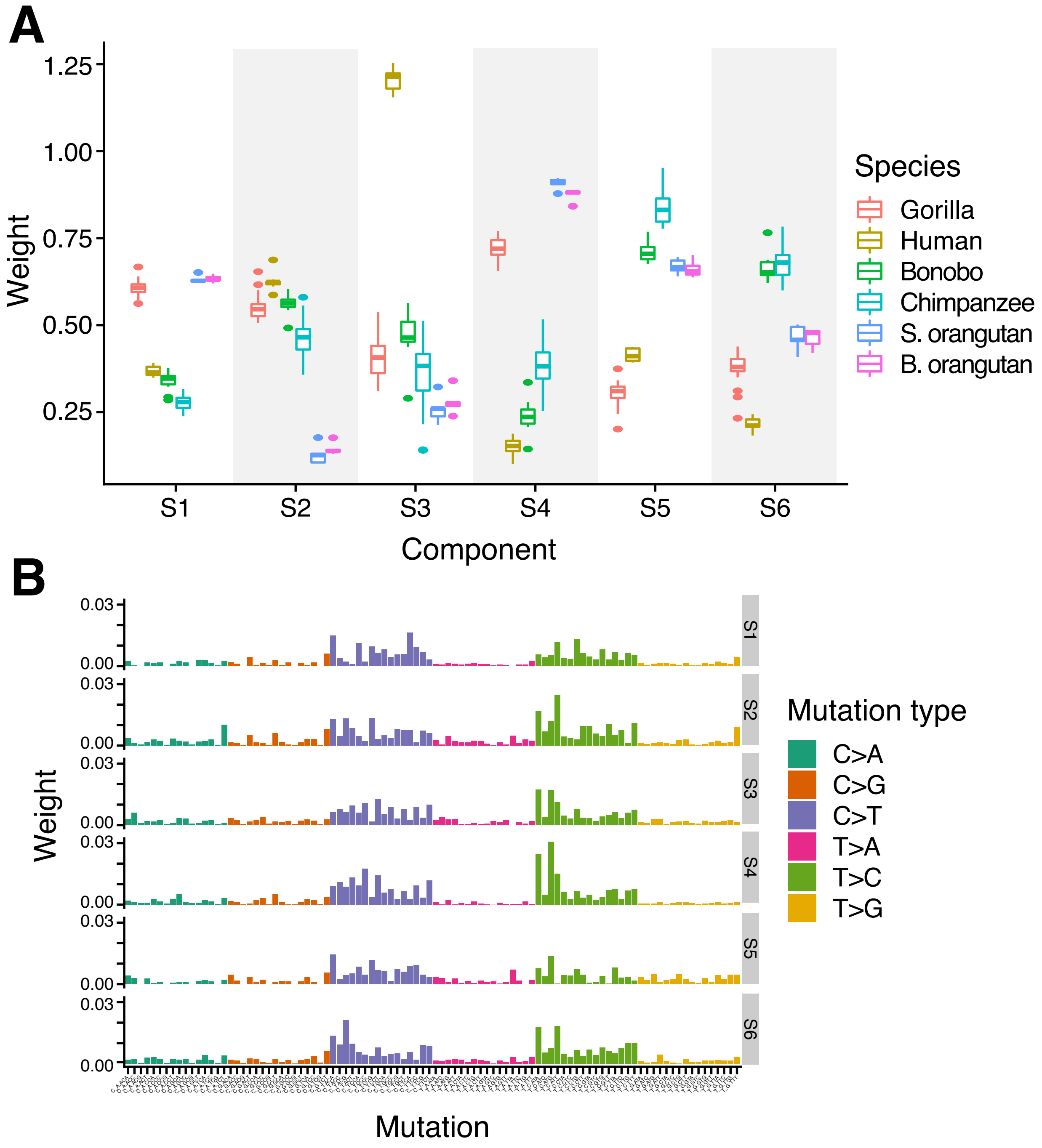


Figure S7: NMF identifies rapidly-evolving germline mutation signatures that are specific to great ape lineages.

A. The dosages of mutation signatures appear lineage-specific. The boxplots shows the distribution of each signature's weight for each species.
B. Different signatures are enriched for different mutation types.

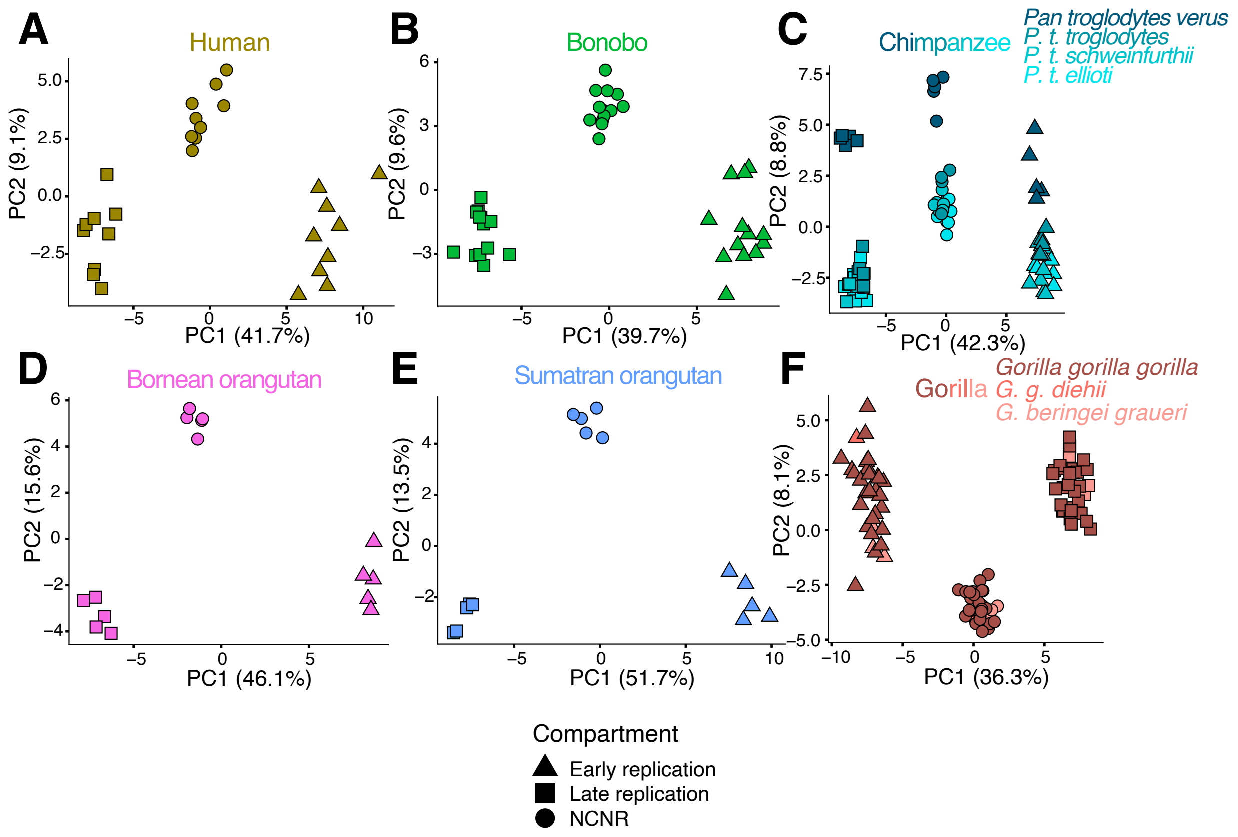


Figure S8
PCA of replication timing and NCNR compartments for individuals in each species. Each point in these PCA represents the mutation spectrum from a single individual's NCNR, early replicating, or late replicating compartment. Clustering by compartment implies differences in mutation spectra based on replication timing.

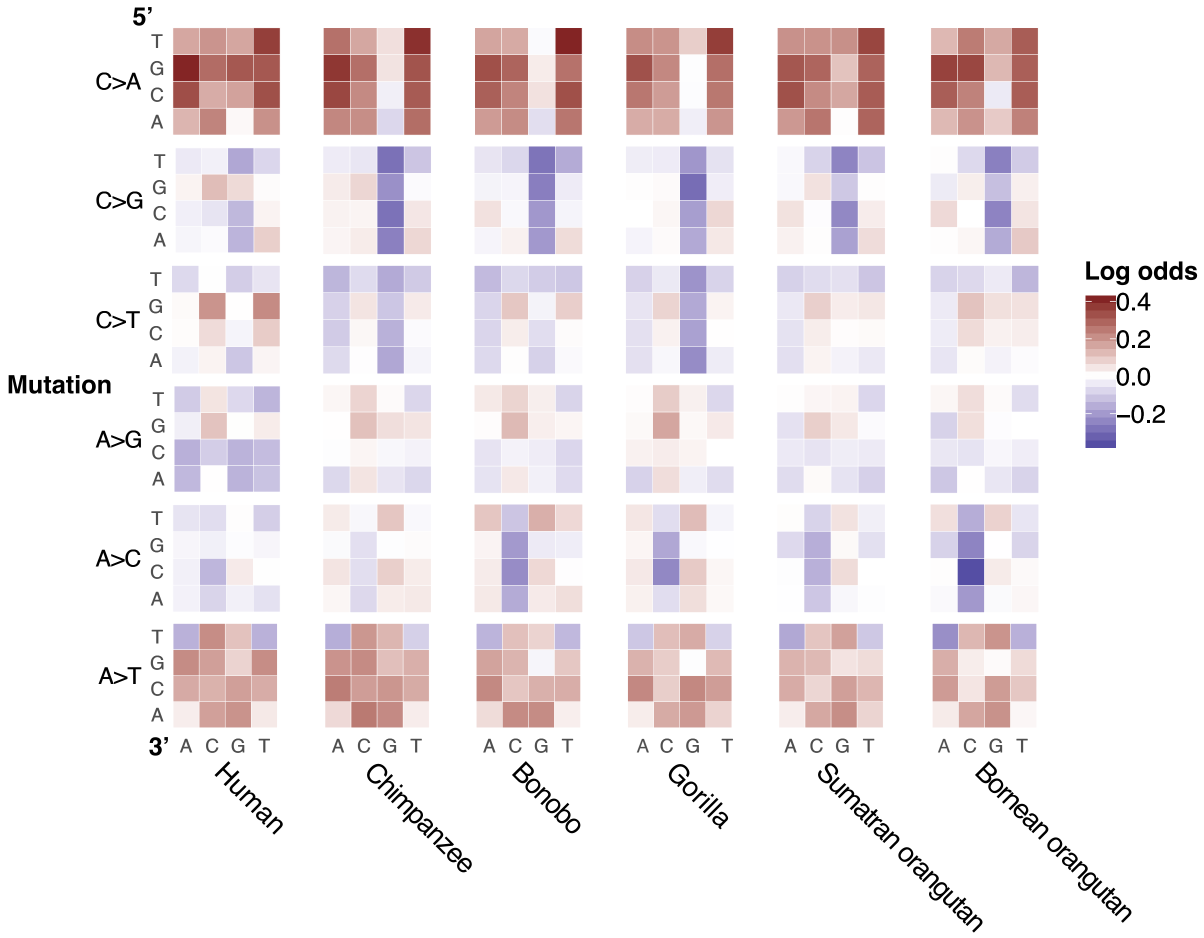


Figure S9
A heatmap of the log ratios of triplet mutation fractions in each species shows an enrichment for C>A and A>T mutations in late replicating compartment compared to early replicating compartment. This mutation signature recapitulates recently described late replication timing signature in humans. To generate the species mutation spectra, we counted the number of SNVs with triplet context segregating within a species that occurred in each compartment. The triplet mutation fractions were normalized by compartment nucleotide content. Statistical quantification of the correlation of these heatmaps can be found in Figure S10.

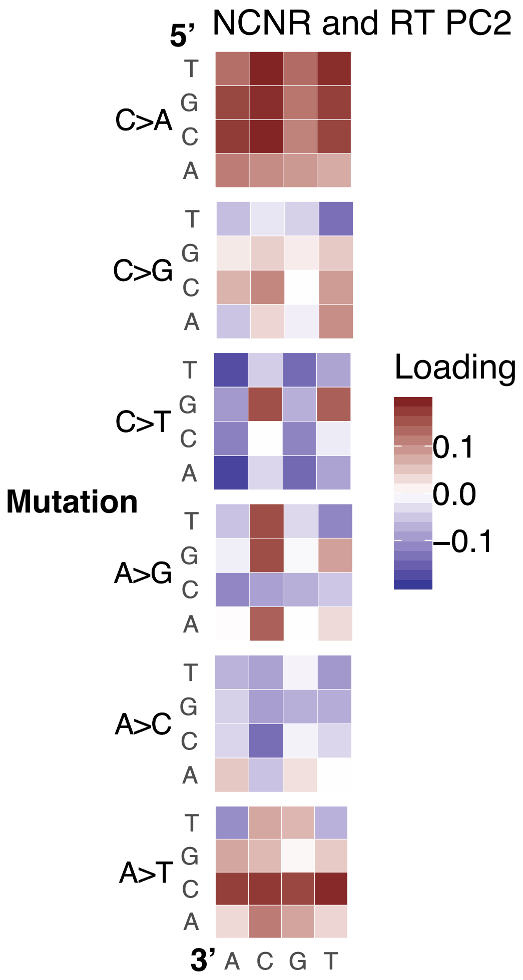


Figure S10

A heatmap showing the PC2 weights associated each triplet mutation type in Figure 2D. The C>A and A>T mutation types dominate PC2, which correlates with a late replication timing signature.

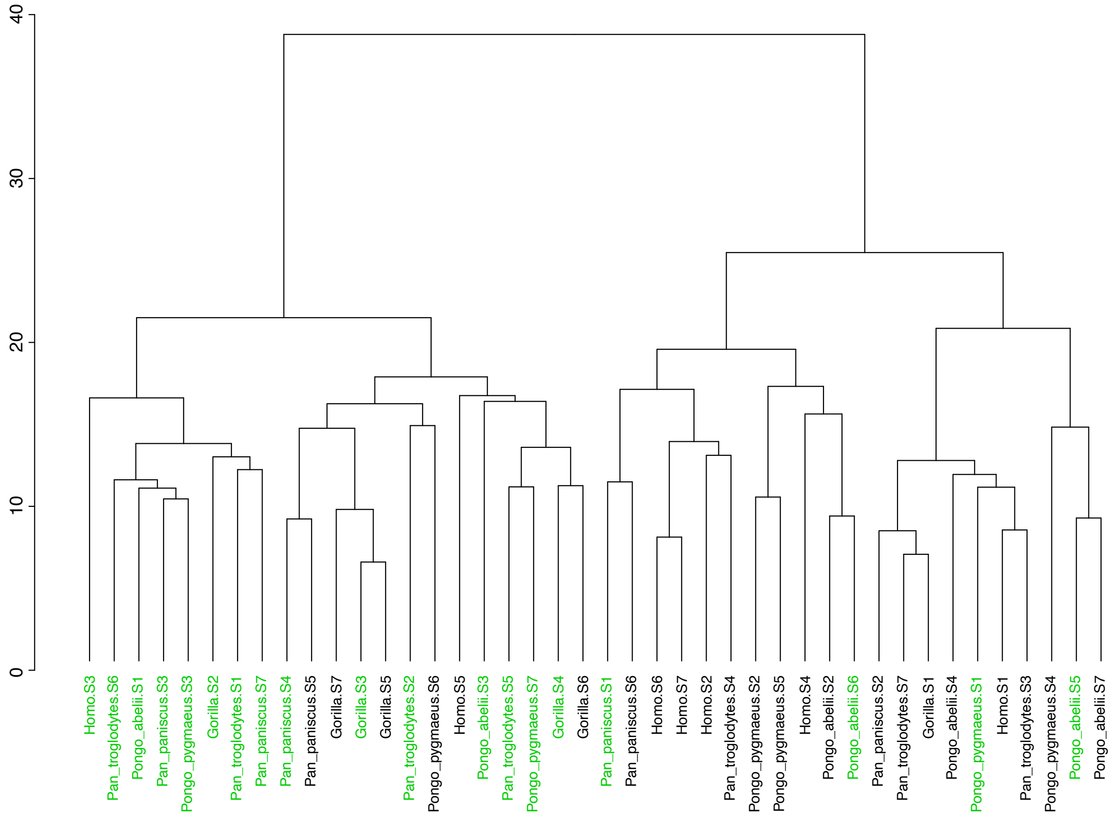


Figure S11. Mutation signatures associated with late-replication timing cluster together. NMF was run on early and late replication timing compartments separately for each species. Distances between these mutation signatures determine the dendrogram's structure; similar signatures are closer together in the tree. Signatures colored in green are enriched in the late replication timing compartment.

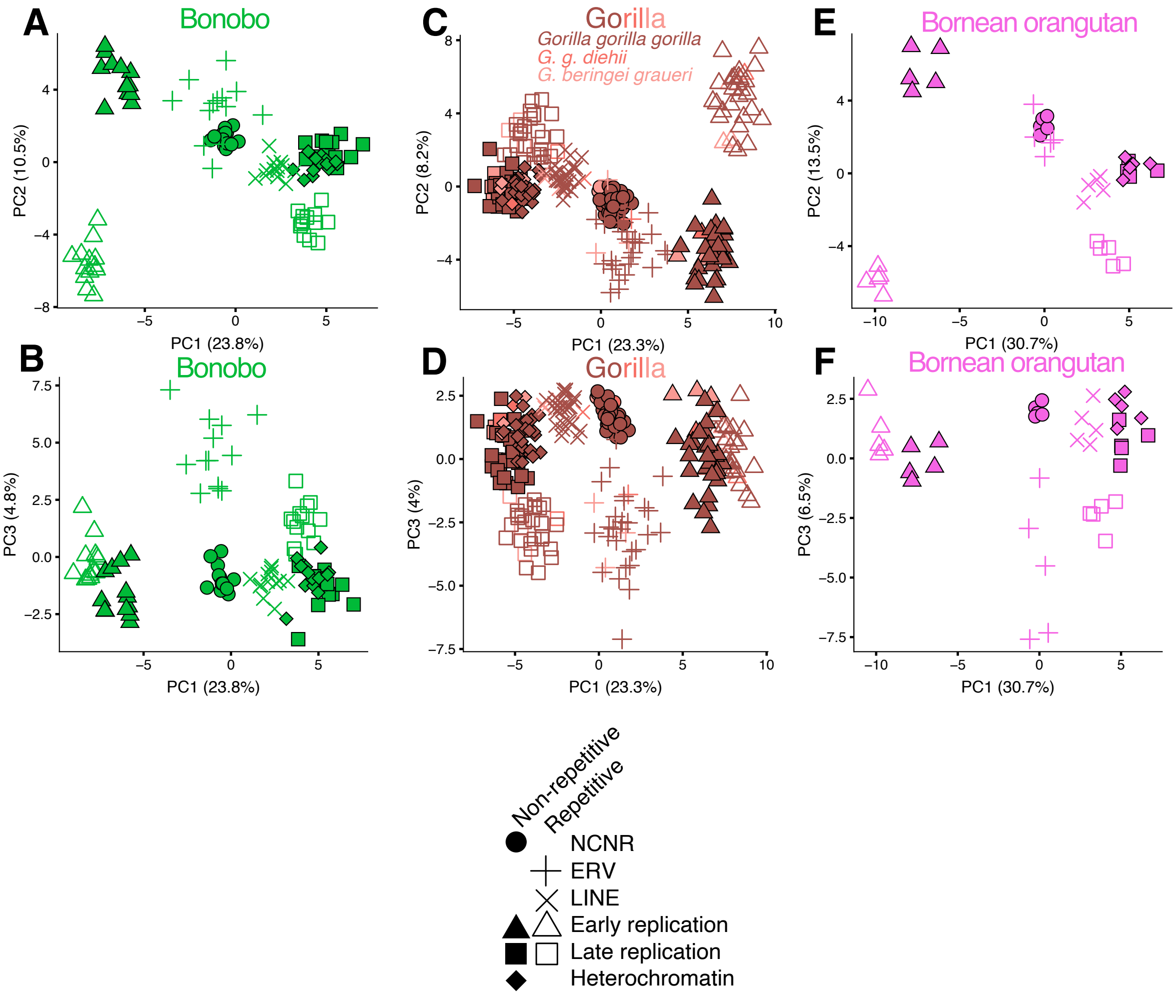


Figure S12
We defined eight overlapping functional compartments to test for evolution of mutation spectrum modifiers along axes of chromatin accessibility, replication timing, and repetitive content. We then ran a PCA on the individual mutation spectra for all eight compartments for each species separately (only bonobo, gorilla, and Bornean orangutan shown here; see Figure 3 for other species). For all species, PC1 and PC2 separate compartments along gradients that correspond to replication timing and repetitive content, respectively. PC3 separates ERVs from other compartments. The similarities of these independent PCAs across all species implies conservation of *cis*-acting mutational signatures.

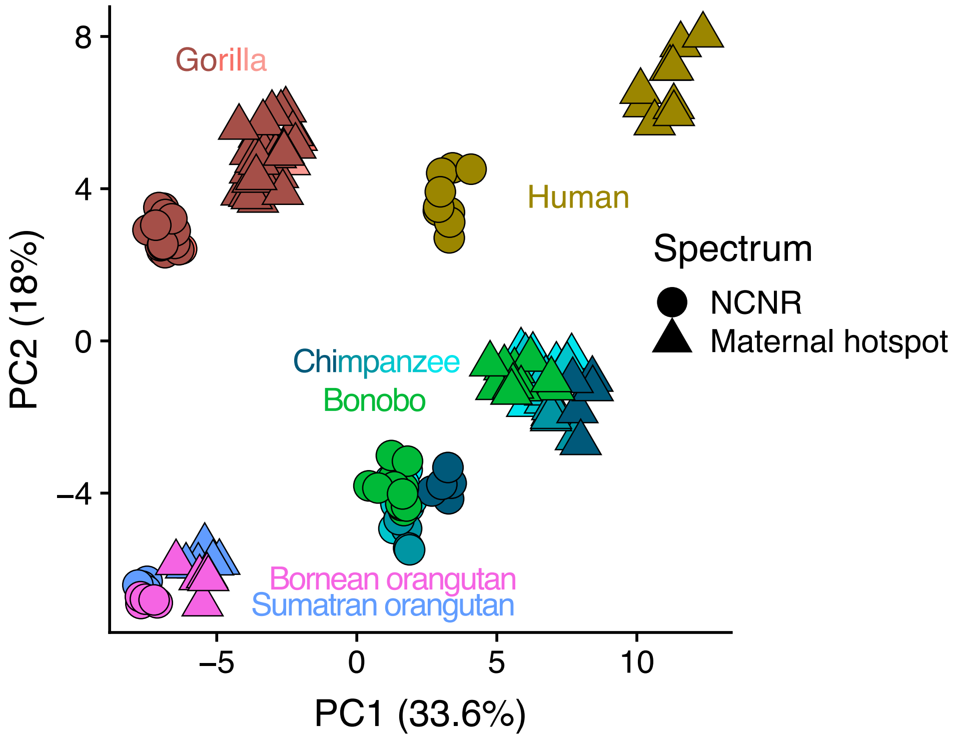


Figure S13
PCA of NCNR and maternal hotspot compartments. Individual mutation spectra for the NCNR and maternal hotspot compartments were calculated and normalized (Methods). The distance between the maternal hotspot and the NCNR mutation spectra is negatively correlated with phylogenetic distance from humans, implying evolution of a *cis*-acting mutational modifier largely absent from orangutans.

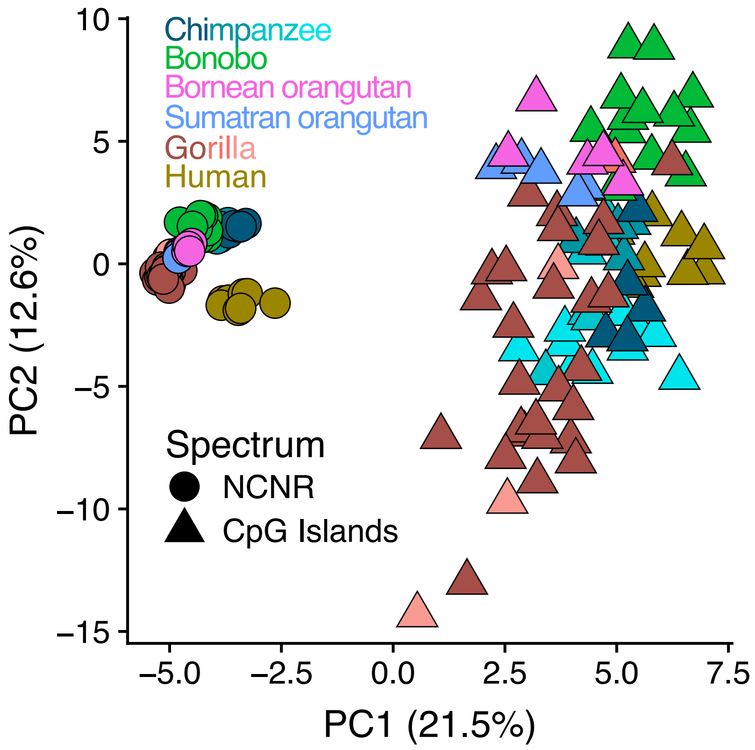


Figure S14
A PCA of NCNR and CpG island compartments demonstrates that differentiation of the CpG island compartment exceeds the magnitude of spectrum differentiation between species.

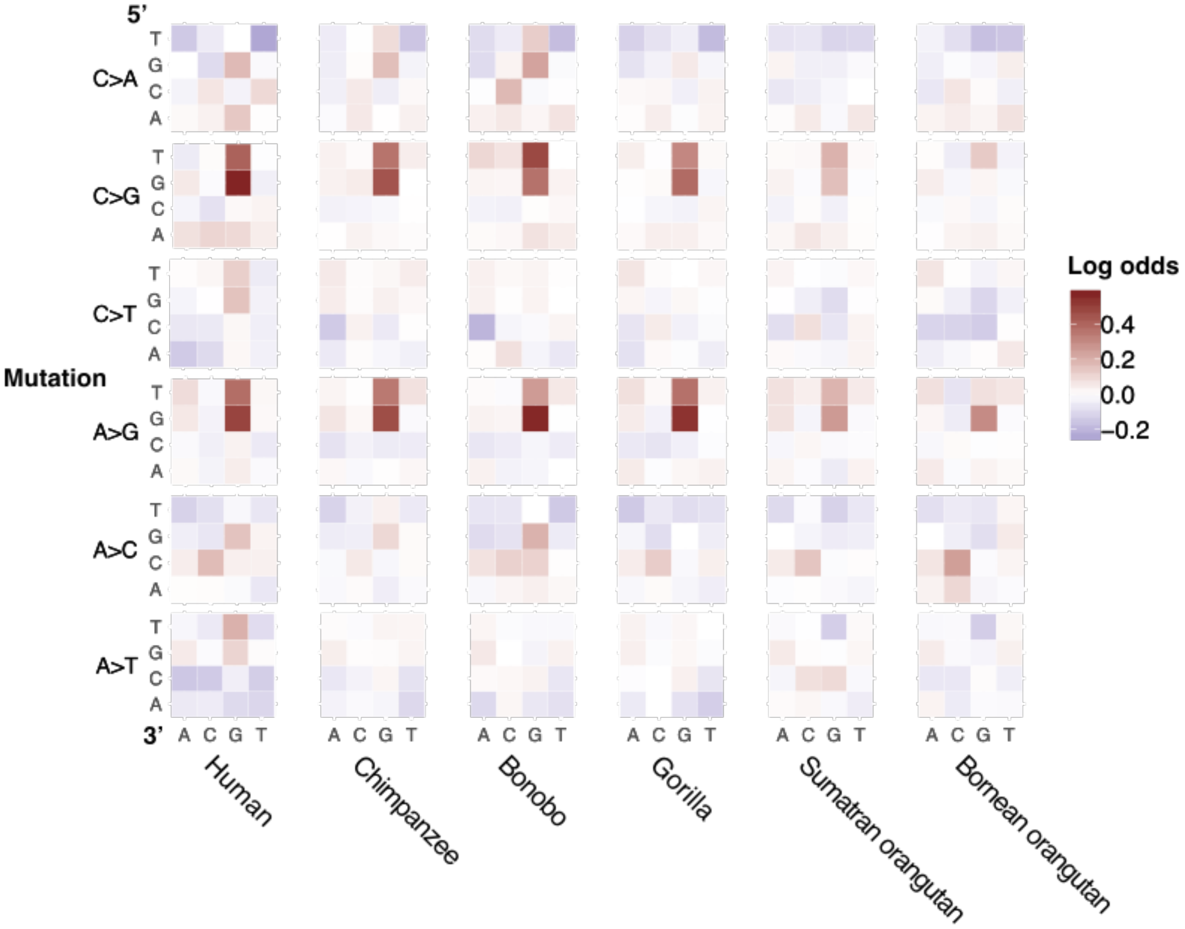


Figure S15
Differences in 7-mer content between the ERV and nonrepetitive heterochromatin compartments do not fully explain the enrichment of CG>GG mutation types in ERVs. We rescaled the counts of each 7-mer mutation type in ERVs by the ratio of the mutating 7-mer's nucleotide content between nonrepetitive heterochromatin and ERV compartments (see Methods). The heatmap shows the log ratio between the 7-mer-corrected 3-mer mutation fractions in ERVs to (uncorrected) 3-mer mutation fractions in nonrepetitive heterochromatin.

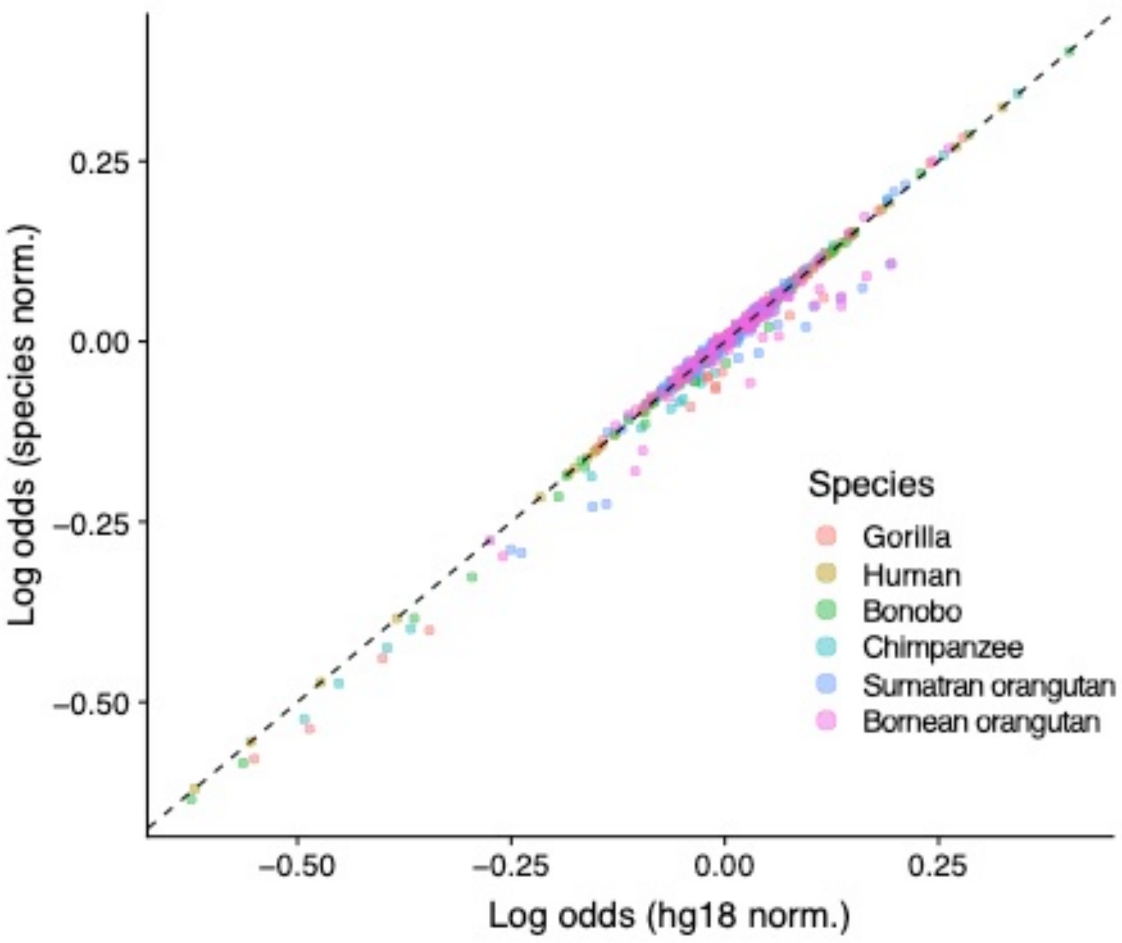


Figure S16
Normalization of mutation spectra is unaffected by variation in species liftovers. We normalized the ERV and nonrepetitive heterochromatin species mutation spectra by the nucleotide content of the genomic regions for each respective compartment the successfully lifted over to each species' reference genome. The log-odds of the ratio of ERV-nonrepetitive heterochromatin mutation spectra are plotted using this species-specific (Y axis) against our standard species-nonspecific normalization (X axis).

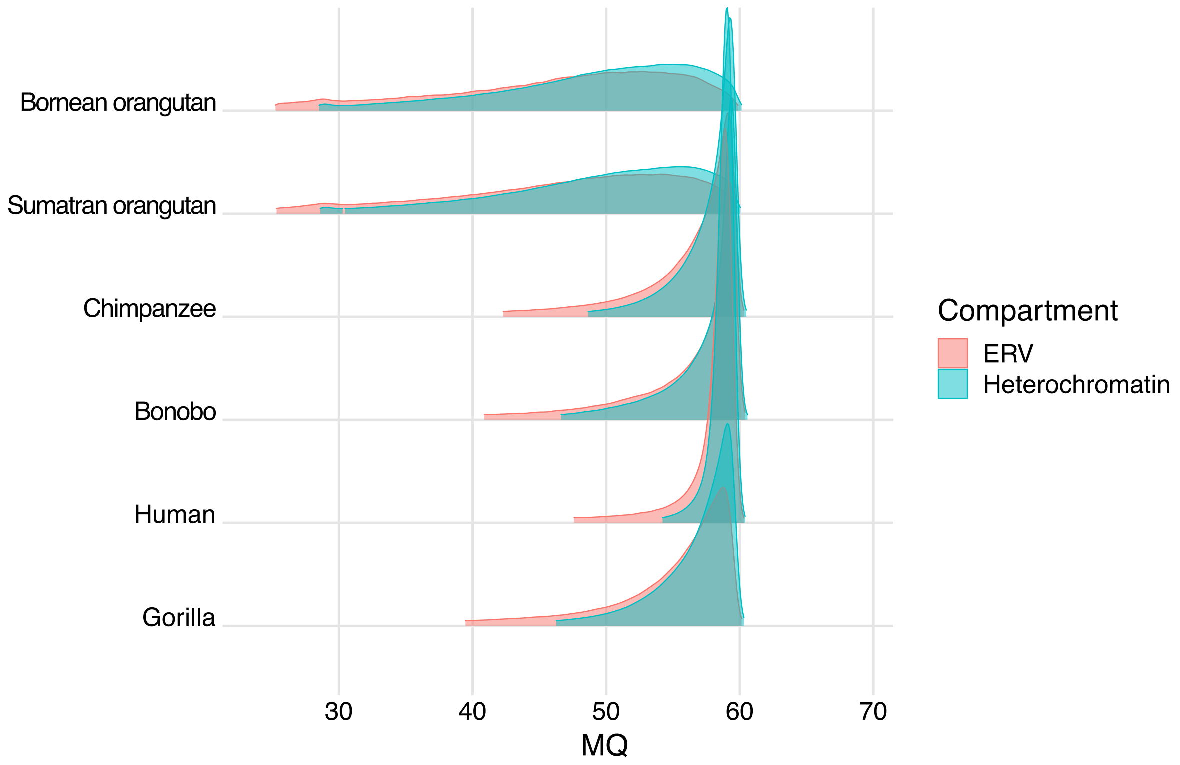


Figure S17
The distribution of SNV mapping quality does not differ substantially between the ERV and nonrepetitive heterochromatin compartments. The plot above shows the layered distributions of mapping quality (MQ in the GAGP VCF files) for SNVs included in the ERV and nonrepetitive heterochromatin compartments in red and blue, respectively. Although each ERV compartment contains a few more low quality SNPs than are present in the matched heterochromatin compartment, they are not numerous enough to explain the spectrum difference between the two regions.

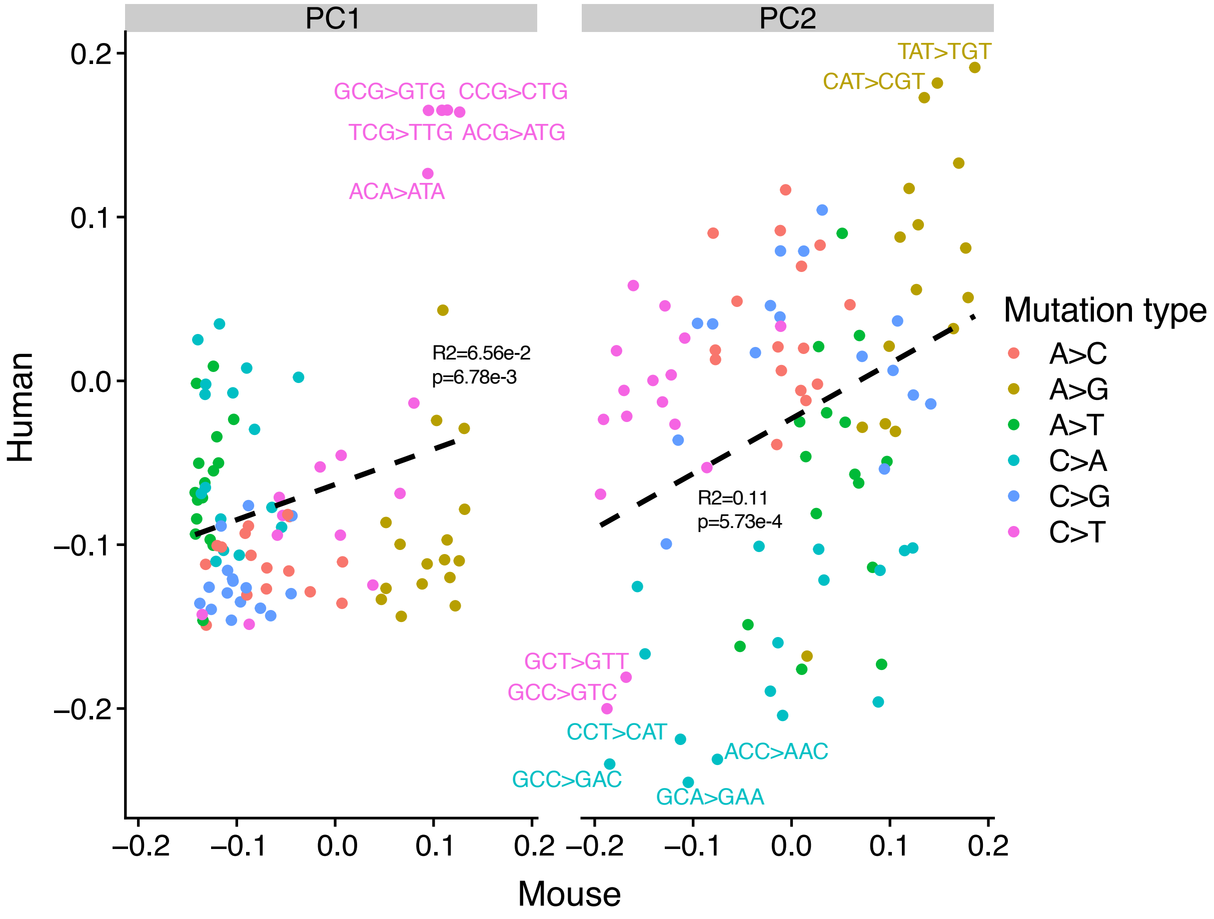


Figure S18

The PC loadings of human and mouse mutation spectra from different functional compartments are significantly correlated. Separate PCAs were run on matrices containing mutation spectra for a set of human and mouse individuals, whose genomes were split into compartments following chromHMM annotations. The annotations were generated for each species separately; thus, regions in the same functional compartment between the two species were not necessarily homologous. The loadings of PC1 and PC2 from the separate PCAs are significantly correlated. Similar mutation types drove the separation and clustering of mutation spectra along both PCs. The dashed line was generated by a linear regression, whose fit and slope significance are listed on the plots.
